# Inhibition of miR-199b-5p reduces pathological alterations in Osteoarthritis by potentially targeting *Fzd6* and *Gcnt2*

**DOI:** 10.1101/2023.10.03.560693

**Authors:** Tong Feng, Qi Zhang, Si-Hui Li, Yan-ling Ping, Mu-qiu Tian, Shuan-hu Zhou, Xin Wang, Jun-Meng Wang, Fan-Rong Liang, Shu-Guang Yu, Qiao-Feng Wu

## Abstract

Osteoarthritis (OA) is a degenerative disease with a high prevalence in the elderly population, but our understanding of its mechanisms remains incomplete. Analysis of serum exosomal small RNA sequencing data from clinical patients and gene expression data from OA patient serum and cartilage obtained from the GEO database revealed a common dysregulated miRNA, miRNA-199b-5p. In vitro cell experiments demonstrated that miRNA-199b-5p inhibits chondrocyte vitality and promotes extracellular matrix degradation. Conversely, inhibition of miRNA-199b-5p under inflammatory conditions exhibited protective effects against damage. Local viral injection of miRNA-199b-5p into mice induced a decrease in pain threshold and OA-like changes. In an OA model, inhibition of miRNA-199b-5p alleviated the pathological progression of OA. Furthermore, bioinformatics analysis and experimental validation identified *Gcnt2* and *Fzd6* as potential target genes of miRNA-199b-5p. Thus, these results indicated that miRNA-199b-5p/*Gcnt2* and *Fzd6* axis might be a novel therapeutic target for the treatment of OA.

## Introduction

Osteoarthritis (OA) is a degenerative disease characterized by the deterioration of articular cartilage and affecting all components of the joint, including the synovium, subchondral bone, and meniscus^1^. Knee OA (KOA) is the leading cause of disability and increased living costs among elderly patients^2^. Currently, the treatment options for osteoarthritis are limited to symptomatic relief and joint replacement. However, the insufficient relief of symptoms, potential medication side effects, and the economic burden and complications associated with joint replacement surgeries have prompted the need to identify new, effective, and safe treatment methods and explore unknown targets for individuals with KOA^3^.

Exosomes are extracellular vesicles that are distributed in body fluids such as serum and play a crucial biological role by delivering molecules such as MicroRNAs (miRNAs)^4^. They serve as vital carriers for intercellular communication and transfer of genetic information. MiRNAs are a class of non-coding RNAs ranging from 18 to 24 nucleotides in length, which exert inhibitory effects on the expression of target genes through interactions with mRNA. Studies have demonstrated that miRNAs play a crucial role as a pathogenic factor in osteoarthritis (OA). ^5^ For instance, miR-199b-5p was found to contribute to the osteogenic differentiation of bone marrow stromal cells ^6^LJanother miR-140 showed cartilage-specific expression, and its expression was significantly reduced in OA cartilage ^7;8^.Recently,intra-articular injection of antisense oligonucleotides of miR-181a-5p produced chondroprotective effects in OA mice ^9^.

In this study, we initially screened miR-199b-5p as a potential key miRNA based on clinical data by detecting differentially expressed exosomal miRNAs. To elucidate the role and function of miR-199b-5p, we investigated its impact on chondrocytes and identified its molecular targets through *in vitro* experiments. Furthermore, we explored its role *in vivo*. Hence, this article not only identifies miR-199b-5p as a potential micro-target for KOA but also provides a potential strategy for future identification of new molecular drugs.

## Results

### Identification and enrichment analysis of differentially expressed miRNAs in serum exosomes

To investigate the dysregulated miRNAs in serum exosomes of KOA patients, we extracted exosomal miRNAs from serum and performed sequencing. Serum samples from 15 patients with KOA and 10 healthy subjects were collected (Supplementary Table 2). After extraction, the serum exosomes were observed under transmission electron microscopy and nanoparticle tracking analysis, revealing a diameter ranging from approximately 70 to 150 nm (Fig. S1A, B). Furthermore, the surface marker proteins CD9, CD63, and CD81 were found to be expressed on the exosomes (Fig. S1C). Next, we performed sequencing of miRNAs in serum exosomes.

The results showed that 88 miRNAs were up-regulated and 89 miRNAs were downregulated in KOA patients compared with the control group based on fold change > 1.5 and *p* < 0.05 (Fig. 1A, B). Afterwards, we performed bioinformatics Gene Ontology (GO) and Kyoto Encyclopedia of Genes and Genomes (KEGG) analyses on these differentially expressed miRNA. Among the top 5 enriched results of up-regulated differentially expressed miRNAs, the CC (Cell GO:0044297, intracellular GO:0005622, and cytoplasm GO:0005737) GO terms were notable. The MF (Translation activator activity GO:0008494, binding to ubiquitin-conjugating enzymes GO:0031624, and possessing transferase activity GO:0016740) GO terms and BP (Cell differentiation GO:0045595 and the stabilization of mRNA through 3’-UTR mediation GO:0070935) GO terms were also significant (Fig. 1C). The top 10 enriched KEGG pathways, PI3K-AKT signaling, Ras signaling, apoptosis, and other types of O-glycan biosynthesis are considered to be of significant importance (Fig. 1D). The analysis of the top 5 down-regulated differential miRNAs reveals interesting findings. the CC (Cell GO:0005623, intracellular GO:0005622) GO terms were particularly noteworthy. The MF (RNA polymerase II transcription factor activity and sequence-specific DNA binding GO:0000981) GO terms, as well as the BP (collagen-activated signaling pathway GO:0038065) GO terms, were also deemed significant (Fig. 1E). Moreover, the analysis of the top 10 enriched KEGG pathways revealed a strong association with the Hedgehog signaling pathway, pluripotency of stem cells, and glycosaminoglycan biosynthesis-keratan sulfate (Fig. 1F).

**Figure 1.**
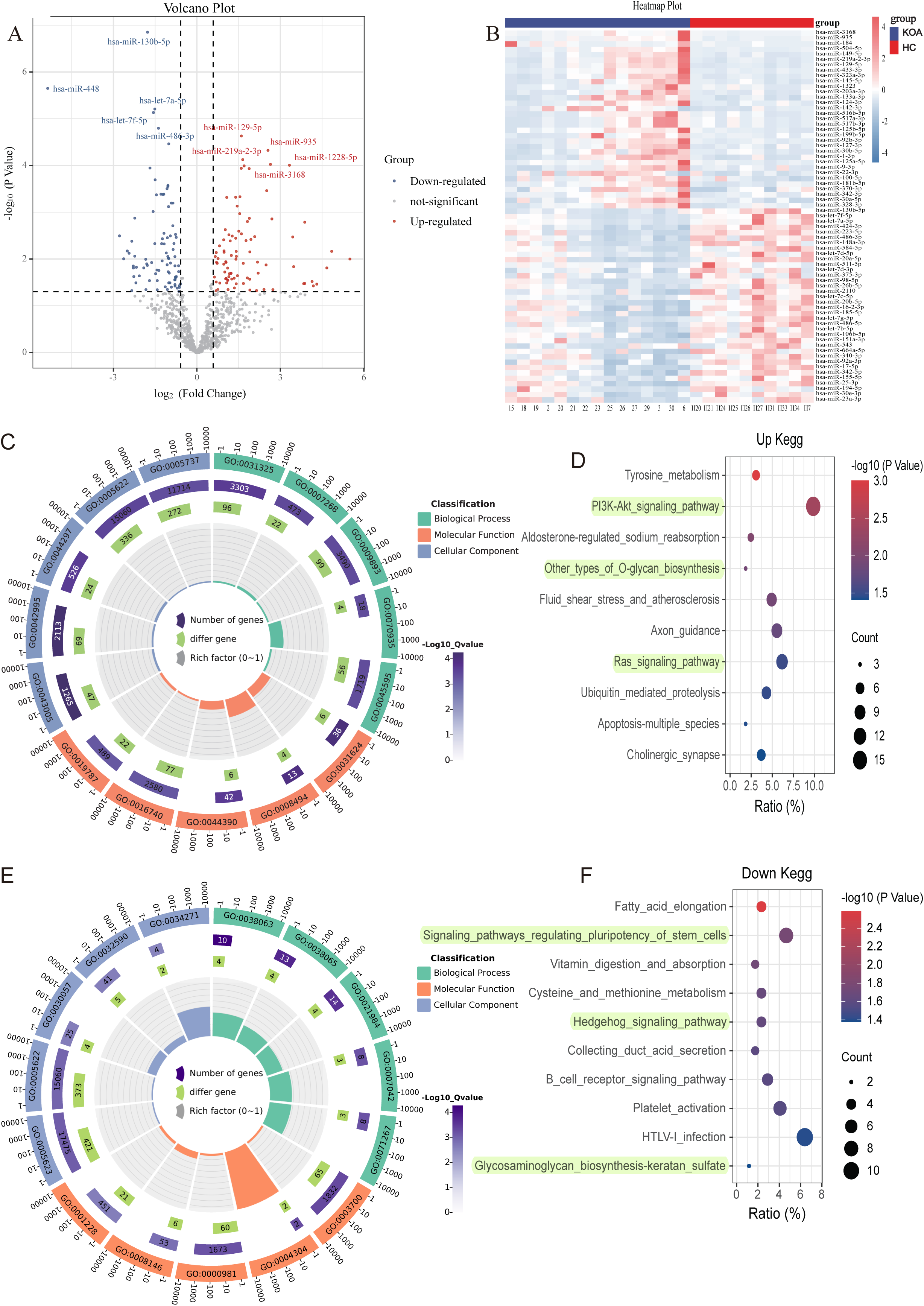
MiRNA expression in KOA patients. **(A, B)** Volcano plot and heatmap showing differential miRNAs between KOA and healthy groups (healthy, *n* = 10; KOA, *n* = 15, ⎢log2 Fold Change⎪≥ 0.585, *P* < 0.05). **(C)** GO enrichment analyses of upregulated genes. (GO:0031325 positive regulation of cellular metabolic process; GO:0007268 chemical synaptic transmission; GO:0009893 positive regulation of metabolic process; GO:0070935 3’-UTR-mediated mRNA stabilization; GO:0045595 regulation of cell differentiation; GO:0043005 neuron projection;GO:0042995 cell projection; GO:0044297 cell body; GO:0005622 intracellular; GO:0005737 cytoplasm; GO:0031624 ubiquitin conjugating enzyme binding; GO:0008494 translation activator activity; GO:0044390 ubiquitin-like protein conjugating enzyme binding; GO:0016740 transferase activity; GO:0019787 ubiquitin-like protein transferase activity) **(D)** KEGG enrichment analyses of upregulated genes. **(E)** GO enrichment analyses of downregulated genes. (GO:0038063 collagen-activated tyrosine kinase receptor signaling pathway; GO:0038065 collagen-activated signaling pathway; GO:0021984 adenohypophysis development; GO:0007042 lysosomal lumen acidification; GO:0071267 L-methionine salvage; GO:0005623 cell; GO:0005622 intracellular; GO:0030057 desmosome; GO:0032590 dendrite membrane; GO:0034271 phosphatidylinositol 3-kinase complex, class III, type I; GO:0003700 DNA binding transcription factor activity; GO:0004304 estrone sulfotransferase activity; GO:0000981 RNA polymerase II transcription factor activity, sequence-specific DNA binding; GO:0008146 sulfotransferase activity; GO:0001228 transcriptional activator activity, RNA polymerase II transcription regulatory region sequence-specific DNA binding) **(F)** KEGG enrichment analyses of downregulated genes.

It is noteworthy that in the aforementioned results, both the up-regulated and down-regulated differential miRNA enrichment analyses are involved in processes related to cartilage synthesis, including O-glycan biosynthesis and glycosaminoglycan biosynthesis-keratan sulfate^10;11^.

### Dysregulated serum exosomal miRNAs were found to be expressed in both serum and cartilage

Subsequently, we aimed to investigate whether these dysregulated miRNAs in serum exosomes show differential expression in other affected sites or tissues of KOA. Leveraging the KOA patient cartilage and serum sequencing data available in the GEO database, we compared our dysregulated miRNA list with the datasets. Remarkably, we identified 169 miRNAs that exhibited differential expression in the serum of KOA patients (Fig. 2A), suggesting their involvement in the disease. Moreover, these miRNAs were also found to be expressed in KOA patient cartilage (Fig. 2B). This robust validation reinforces the reliability of our data. Consequently, we proceeded with further screening of differentially expressed genes.

**Figure 2.**
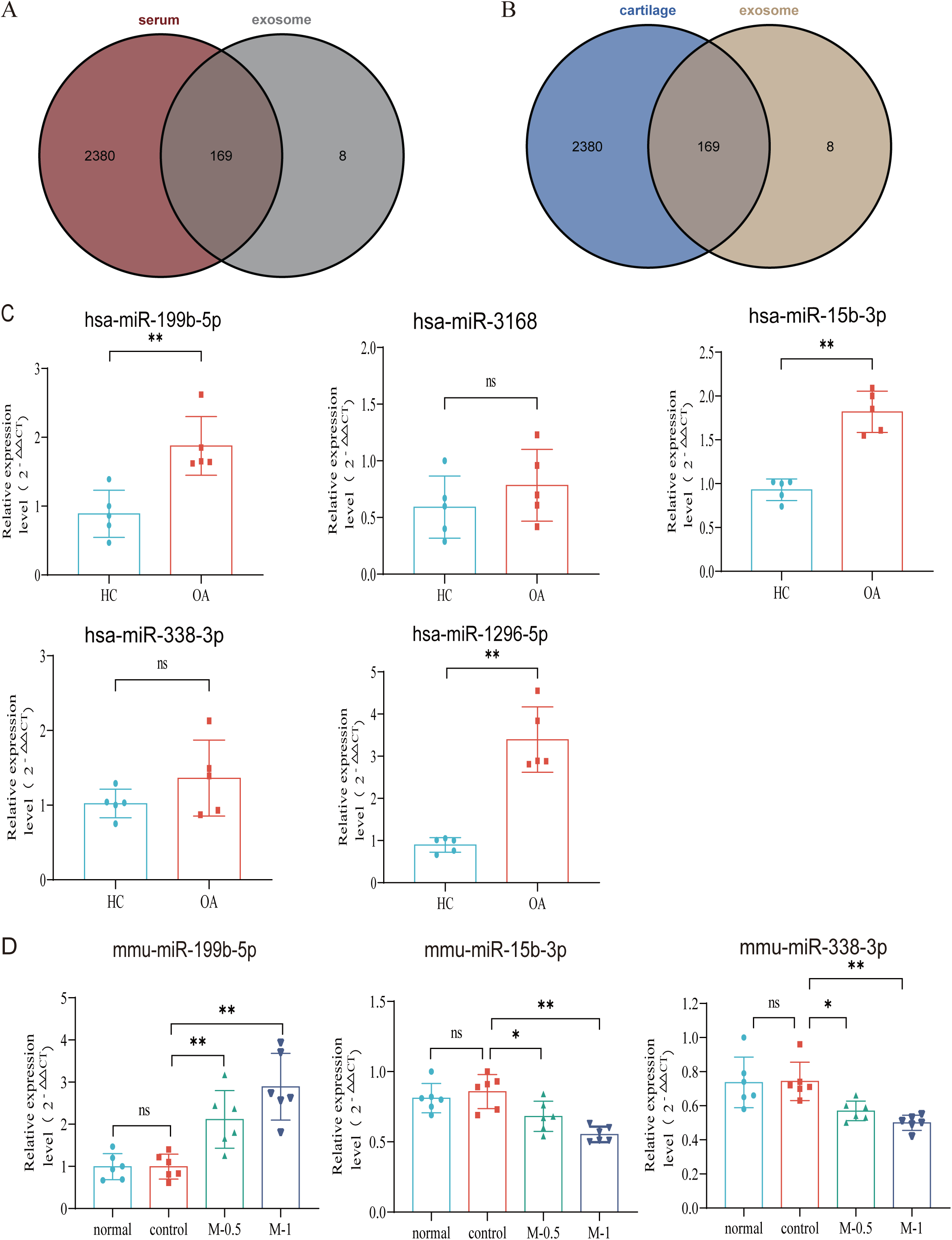
Verification with GEO data and miRNA screening. **(A, B)** Venn plot showing GEO dataset and our result. **(C)** RT**-**qPCR in clinical samples to verify the expression (HC, *n* = 5, KOA, n = 5). **(D)** RT**-**qPCR results of mouse joint samples (*n* = 6). M-0.5, 10 µL of 0.5 mg/mL;M-1, 10 µL of 1 mg/mL. Data are shown as means ± SD. \**P*<0.05, \*\**P*<0.01.

### miRNA-199b-5p has been identified as a potential target molecule in KOA

Based on the P-value and exosomal expression, combined with the results of our additional experiments (Fig. S5). Five miRNAs (hsa-mir-3168, hsa-mir-1296-5p, hsa-mir-15b-3p, hsa-mir-338-3p and hsa-mir-199b-5p) were selected to further research and validated in independent human samples by RT-qPCR (hsa-mir-199b-5p, *p* < 0.01; hsa-mir-3168, *p*=0.33; hsa-mir-15b-3p, *p* < 0.01; hsa-mir-338-3p, *p*=0.20 and hsa-mir-1296-5p, *p* < 0.01) (Fig. 2C). To further explore the functional roles of these miRNAs, we established a mouse model of KOA to evaluate their expression in the joints. Due to species differences, the corresponding sequence of mir-1296-5p could not be identified in the mouse model. we examined the expression of miR-199b-5p (*p* <0.01), miR-15b-3p (*p* <0.05) and miR-338-3p (*p* <0.05) in mouse joint tissue samples (control *vs.* M-0.5) (Fig. 2D). The results showed that only the expression trend of miR-199b-5p was consistent between the clinical samples and the mouse arthritis model. Therefore, we selected miR-199b-5p as the target for our subsequent research.

### miR-199b-5p mimic inhibits the cell viability and extracellular matrix (ECM) of chondrocytes while the inhibitor restores LPS-induced chondrocytes damage

We first extracted mouse primary chondrocytes (Fig. 3A-C). In vitro experiments showed that overexpression miR-199b-5p inhibited the viability of chondrocytes(*p*<*0.01*) (Fig. 3D, Fig. S2). we also find miR-199b-5p mimic increased the mRNA expressions of *MMP3* (*p=*0.09) and *ADAMTS5* (*p<0.05*) and decreased the mRNA expression of *COL2A1*(*p*=0.05), *AGGRECAN* (p=0.20) and *SOX9* (*p*=0.22), which are often used as the biomarkers of chondrocytes ECM metabolic balance. In contrast, miR-199b-5p inhibitor decreased the mRNA expression of *MMP3* (*p*<0.01) and *ADAMTS5* (*p*=0.11) and increased the mRNA expression of *COL2A1* (*p*=0.07), *AGGRECAN* (*p*<0.01) and *SOX9* (*p*<0.01) (Fig. 3E–I). While some gene expression changes may not be significant in statistic, but the modulation of miR-199b-5p expression has been observed to exert an influence on the metabolic alterations of chondrocytes.

**Figure 3.**
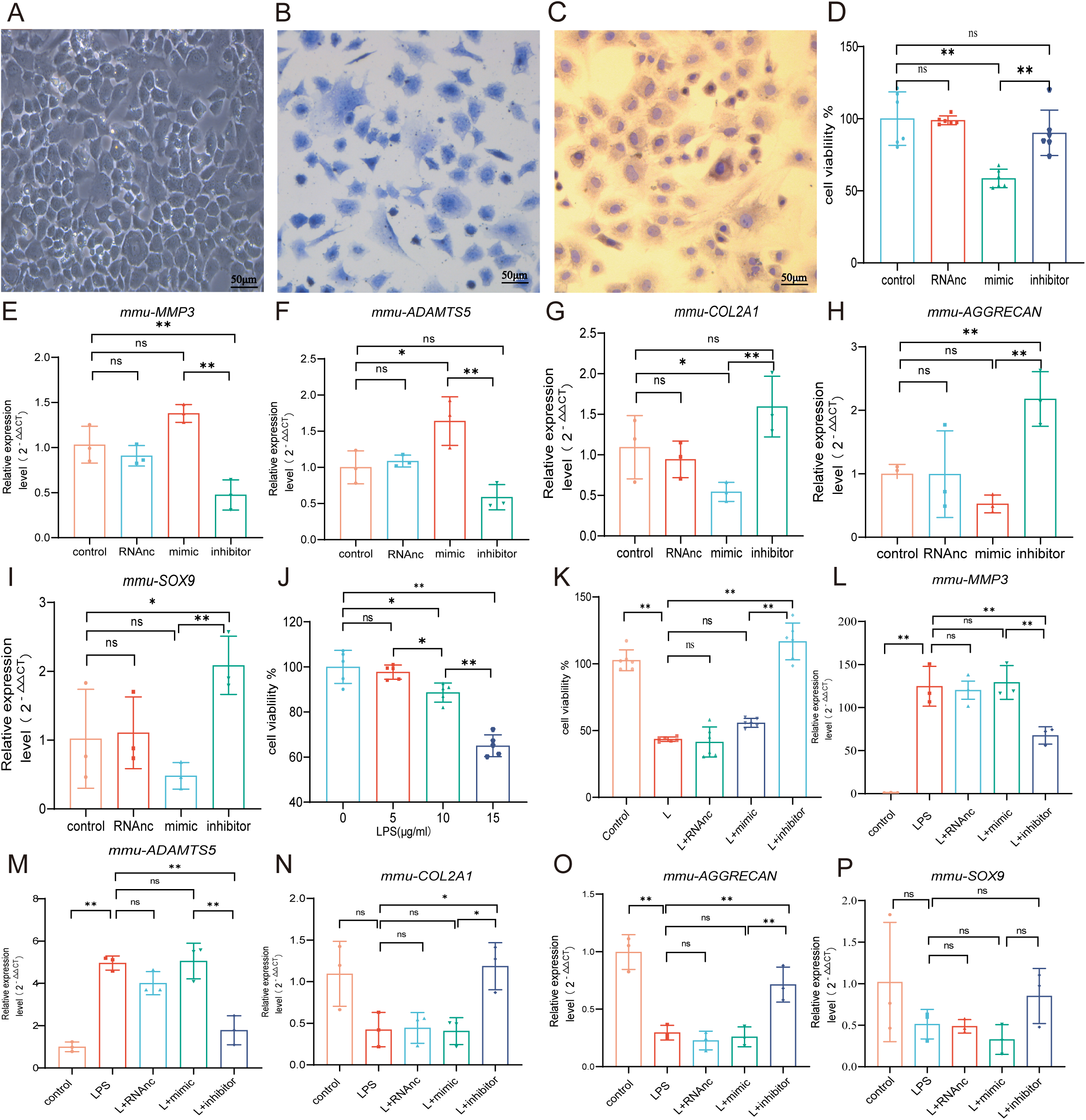
Chondrocyte proliferation and marker expression changes after miR-199b-5p mimic or inhibitor treatment. **(A)** Second generation primary mouse chondrocytes. **(B)** Toluene blue staining. **(C)** Type LJ collagen immunoassay. **(D)** CCK-8 assay for cell viability (*n*=6). **(E, F)** RT-qPCR detection of *MMP-3* and *ADAMTS5* mRNA expression (*n*=3). **(G-I)** RT**-**qPCR detection of *COL-2A1*, *AGGRECAN*, and *SOX9* mRNA expression (*n*=3). **(J)** CCK-8 cell viability assay after different doses of LPS induction (*n*=5). **(K)** CCK-8 cell viability assay after virus infection (*n*=6). **(L, M)** RT**-**qPCR detection of *MMP-3* and *ADAMTS5* mRNA expression (*n*=3). **(N-P)** RT**-**qPCR detection of *COL2A1*, *AGGRECAN*, and *SOX9* mRNA expression (*n*=3). Data are shown as means ± SD. \**P*<0.05, \*\**P*<0.01.

To explore the effect of miR-199b-5p under pathological conditions, an LPS-stimulated inflammation chondrocyte cell model was established. We examined the effect of LPS at 5, 10, and 15 μg/ml on cell viability and found that cell viability was significantly decreased at 15 μg/ml(*p*<0.001) (Fig. 2J). Next, we chose 15 μg/ml of LPS to establish a chondrocyte injury model. The CCK-8 assay showed that miR-199b-5p inhibitor reversed the decrease of cell viability caused by LPS (*p*<0.01) (Fig. 3K). Also, we revealed that LPS elevated *MMP3* (*p*<0.01) and *ADAMTS5* (*p*<0.01) and decreased *COL2A1*(*p*=0.06), *AGGRECAN* (*p*<0.01), and *SOX9* (*p*=0.13) expression. In the presence of miR-199b-5p inhibitor, the changes in mRNA levels were reversed (Fig. 3L-P). These results suggest that miR-199b-5p overexpression reduces cell viability and miR-199b-5p inhibition partly restores LPS-induced cell damage and ECM degradation.

### miR-199b-5p mimic induces inflammation in normal mice, and miR-199b-5p inhibitor alleviates symptoms in KOA mice

Now, we want to know the vivo role of miRNA-199b-5p. Firstly, we screened Adneovirus (AD) and utilized High-AD as a vector to either overexpress or inhibit the expression of miR-199b-5p(Fig. S3). In normal mice, miR-199b-5p mimic was injected to the joint of mice (Fig. 4A) and a decrease in pain threshold was found (*p*<0.01) (Fig. 4B). Four weeks later, serum *IFN-*_γ_ (*p*<0.01) and *TNF-*_α_ (*p*<0.01) were also significantly increased (Fig. 4C, D). The injection of miR-199b-5p mimic in mouse joints resulted in a slight decrease in Safranine staining, although it did not reach statistical significance (Fig. 4E, F). Additionally, the articular surface μCT image showed slight erosion (Fig. 4G). interestingly, the level of serum inflammation in the miR-199b-5p inhibitor injection group was significantly decreased compared with the mimic group (*IFN-*_γ_, *p*<0.01; *TNF-*_α_, *p*<0.01). These results indicated that intra-articular injection of miR-199b-5p mimic induced inflammation response in normal mice.

**Figure 4.**
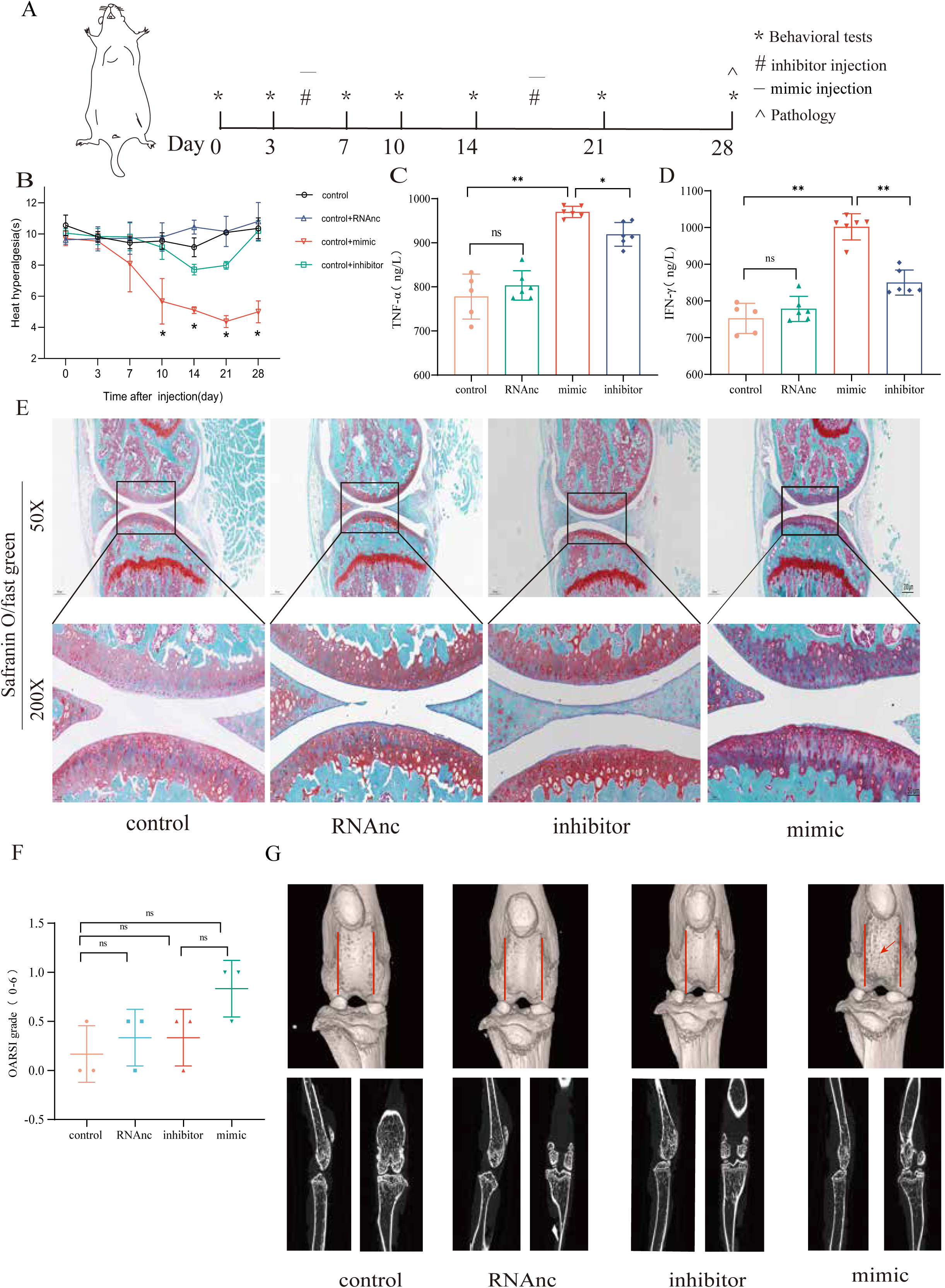
Injection of adenovirus expressing miR-199b-5p mimic results in inflammation and pain threshold sensitivity in mice. **(A)** Animal experiment schematic. **(B)** Behavioral detection of animal thermal pain threshold. **(C, D)** Detection of serum levels of *IFN-*_γ_ and *TNF-*_α_ in controls by ELISA. **(E, F)** Safranin-fast green staining and semiquantitative scoring of articular cartilage. **(G)** 3D reconstruction and 2D images of joints from μCT scans. Data are shown as mean ± SD. \**P*<0.05, \*\**P*<0.01, *n*=6.

Followingly, we established an MIA-induced KOA model to further investigate the role of miR-199b-5p (Fig. S4). We observed a decrease in pain threshold in the model group, and recovery was observed on the 10th day in the inhibitor group (*p*<0.01) (Fig. 5A, B). Joint tissues were taken at the fourth week, and revealed that the expression of *IFN-*_γ_ (*p*<0.01) and *TNF-*_α_ (*p*<0.01) decreased after inhibiting miRNA-199b-5p (Fig. 5C, D). Safranin-fast green staining of joints showed recovery of articular cartilage degradation (*p*<0.05) (Fig. 5E, F), μCT image demonstrated a partial amelioration of the cartilage erosion. (Fig. 5G). These results proved that intra-articular injection of the miR-199b-5p inhibitor partly recovered pain, inflammation, and cartilage degeneration in KOA mice.

**Figure 5.**
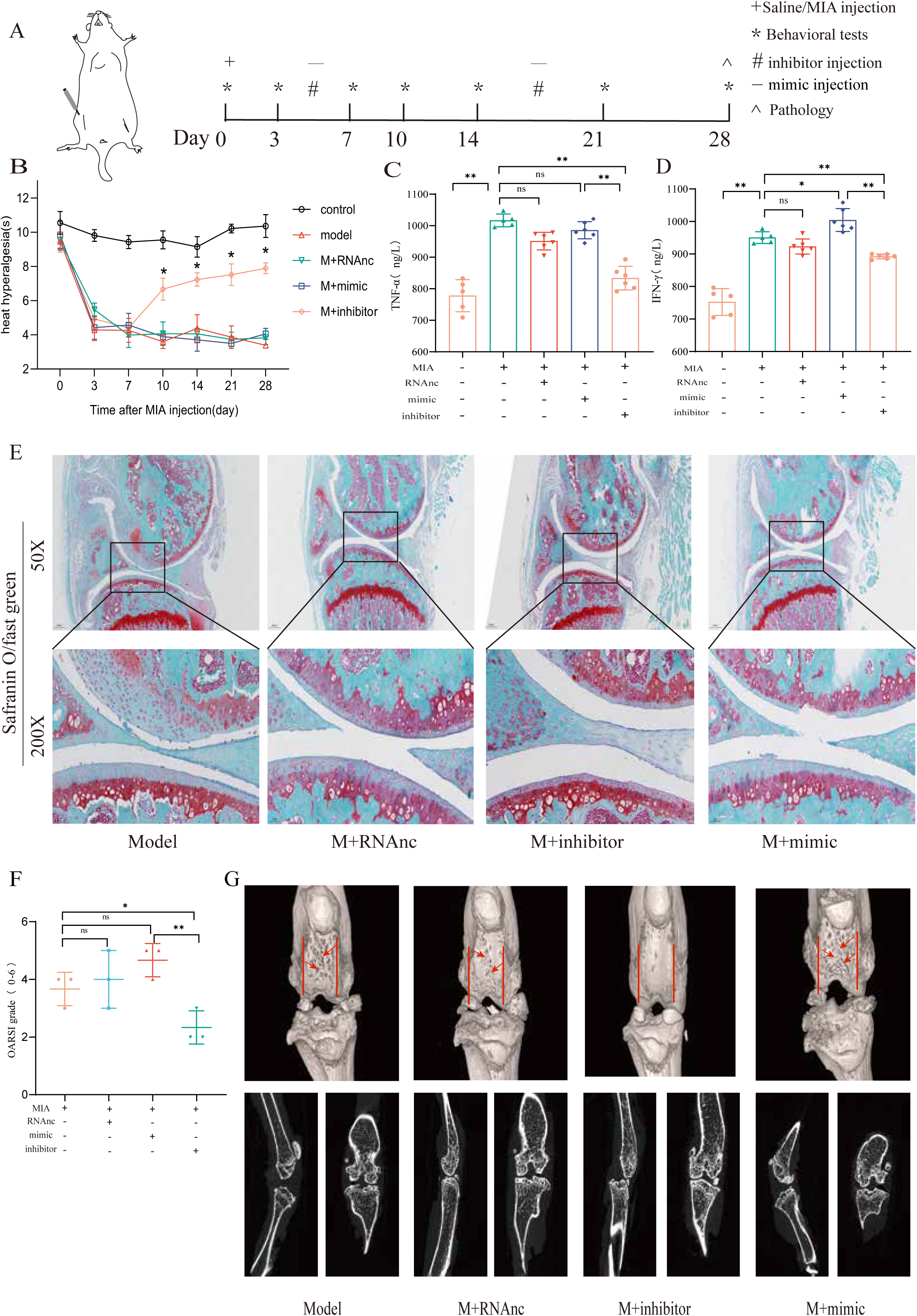
Injection of adenovirus expressing miR-199b-5p inhibitor partly recovered pathological changes in KOA mice. **(A)** Animal experiment schematic. **(B)** Behavioral detection of animal thermal pain threshold. **(C, D)** Detection of serum levels of *IFN-*_γ_ and *TNF-*_α_ in controls by ELISA. **(E, F)** Safranin-fast green staining and semiquantitative scoring of articular cartilage. **(G)** 3D reconstruction and 2D images of joints from μCT scans. Data are shown as mean ± SD. \**P*<0.05, \*\**P*<0.01, *n*=6.

### *Gcnt2* and *Fzd6* are two potential target genes of miR-199b-5p

In order to investigate the underlying mechanism of miRNA-199b-5p, we utilized five widely used miRNA target gene prediction tools, namely miRWalk, miRDB, TarBase, starbase, and TargetScan, to identify potential target genes. Consequently, we identified six putative target genes of miRNA-199b-5p (Fig. 6A). Bioinformatics analysis of the six possible target genes showed that BP is in posttranscriptional regulation of gene expression and angiogenesis; CC is in cytoplasmic vesicle and cell leading edge; and MF is in ubiquitin protein ligase binding and protein domain specific binding (Fig. 6B).

**Figure 6.**
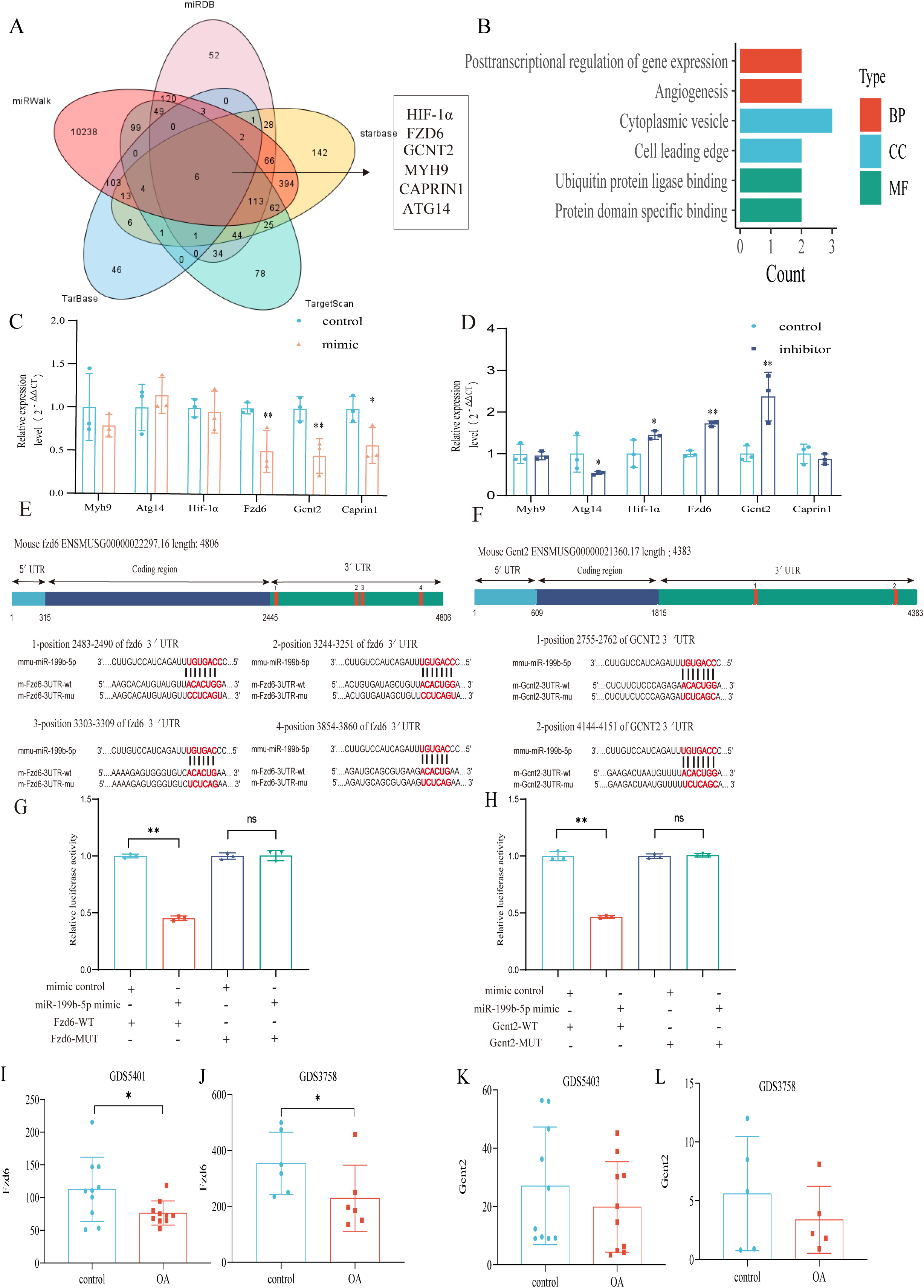
Validation of the miR-199b-5p target gene. **(A)** Prediction of target genes of miR-199b-5p using search sites. **(B)** Target gene GO analysis. **(C, D)** Detection of the expression of target genes under different conditions (*n*=3). **(E, F)** we use targetscan to predict the binding site of miR-199b-5p and target genes. **(G, H)** Validation by luciferase reporter gene assay (n=3). Data are show as mean ± SD.\**P*<0.05, \*\**P*<0.01, *n*=6. **(I-L)** The expression of FZD6 and GCNT2 in the synovial membrane and chondrocytes of GEO Profiles KOA.

After Mimic infected cells, we found decreases expression in *Fzd6* (P<0.01), *Gcnt2* (P<0.01), and *Caprin1* (P=0.024) in the chondrocytes (Fig. 6C). Conversely, after inhibitor infection, *Hif-1*_α_ (P=0.048), *Fzd6* (P<0.01) and *Gcnt2* (P<0.01) were increased and *Atg14* (P=0.042) increased (Fig. 6D). Notably, we observed corresponding changes in the expression of *Fzd6* and *Gcnt2* upon miR-199b-5p overexpression and under-expression. Moreover, we predicted potential binding sites for miRNA-199b-5p within the 3’-untranslated region (UTR) of these two target genes (Fig. 6E, F) and luciferase reporter assays confirmed that miR-199b-5p can potentially bind to and suppress the expression of both *Gcnt2* (P < 0.01) and *Fzd6* (P < 0.01) *via* their complementary sequences (Fig. 6G, H). Additionally, we carried out the comparative analysis of sequence conservatism between human and mouse, and find the binding site on 3’UTR matches to human sequence very well. Besides being positionally conserved, the sequence conservation between hsa_miR-199b-5p and mmu_miR-199b-5p was as high as 95.65%, indicating a good sequence conservation (Fig. S6).

Furthermore, we found differential expression of *Fzd6* in both synovial tissue data (GDS5401) and chondrocyte data (GDS3758) from KOA patients in the GEO profile (Fig. 6I, J). Similarly, differential expression tendency of *Gcnt2* was observed in both the synovial tissue data (GDS5403) and GDS3758 from the same KOA patients (Fig. 6K, L). These findings provide further validation to our results. Therefore, *Fzd6* and *Gcnt2* was considered by us as an important downstream target for further research.

## Discussion

In this study, we initially performed sequencing of serum exosomal miRNAs from clinical patients and identified 177 dysregulated miRNAs. Subsequently, through comparison with GEO data, we found that 169 miRNAs were expressed in both KOA serum and cartilage. Following a screening process, miR-199b-5p was selected for further experiments. In cell-based assays, we discovered that miR-199b-5p can influence the viability of chondrocytes and cytokine-mediated extracellular matrix metabolism. Moreover, in vitro experiments demonstrated that it can induce inflammation and abnormal pain threshold in normal mice. Importantly, inhibition of miR-199b-5p alleviated the pathological symptoms of KOA. Finally, these effects were achieved by potential targeting *Gcnt2* and *Fzd6* (Fig. 7). Thus, our findings demonstrated that miR-199b-5p might be a novel potential therapeutic target for OA prevention and treatment.

**Figure 7.**
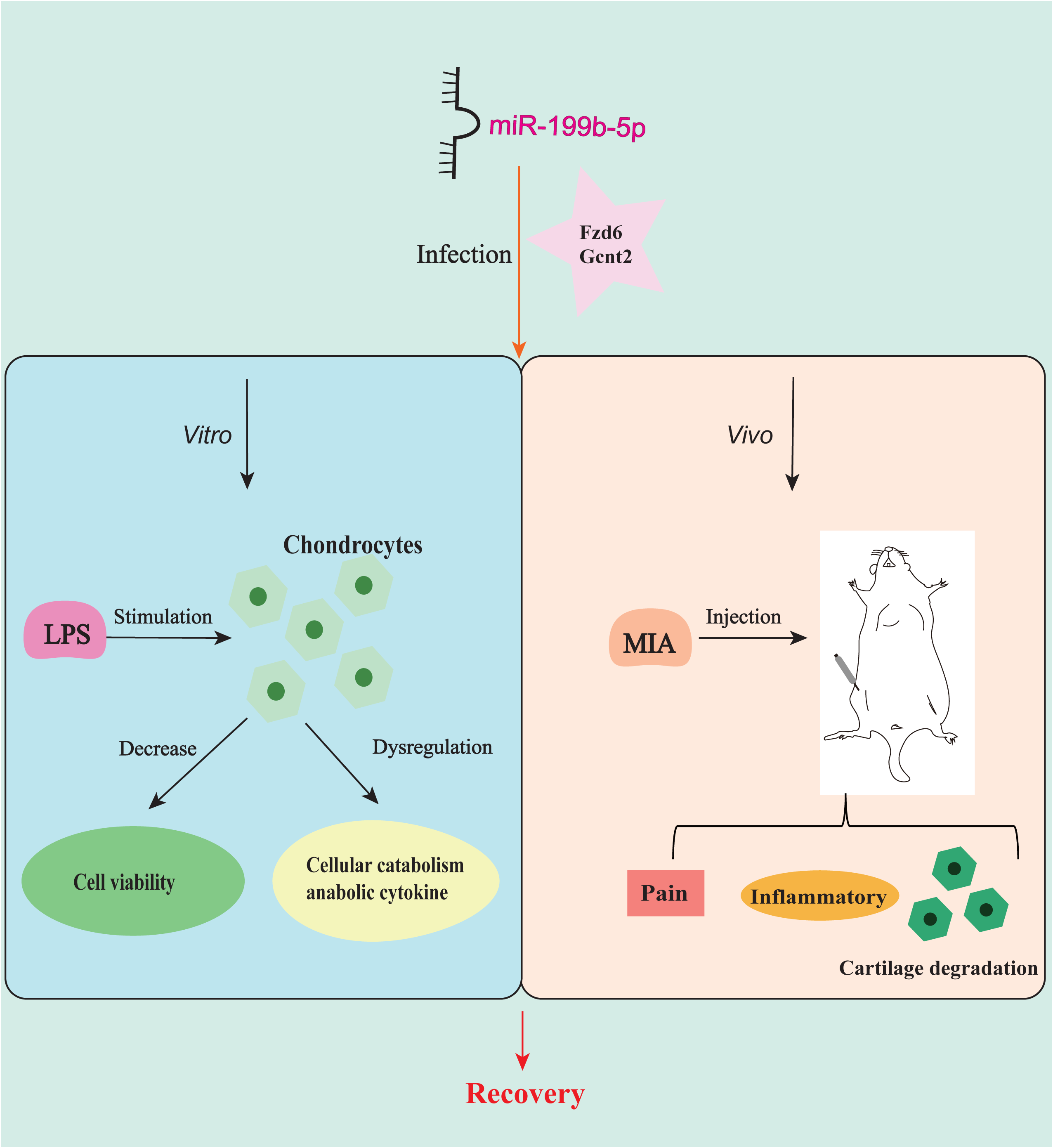
miR-199b-5p exerts its effects on *in vitro* cells and *in vivo* mice by potential targeting *Fzd6* and *Gcnt2*.

miRNAs function as regulators of gene expression in biological processes by regulate mRNA translation by specifically binding to the 3′UTRs of target mRNAs ^12^. Exosomes are rich in miRNAs, and various cells can secrete exosomes to target cells under physiological and pathological conditions, which function as delivery vehicles for miRNAs ^13;14^. There is evidence suggesting that miRNAs can participate in various cellular processes (inflammation, cell viability, ECM dysregulation) and signaling pathways (Hedgehog signaling, PI3K-AKT signaling) relevant to OA^15;16^. Our differential miRNA enrichment analysis also strongly supports the correlation with these findings.

In cancer research, overexpression of miR-199b-5p can inhibit the proliferation, migration and invasion of prostate cancer cells in vitro and tumor growth and metastasis in vivo by targeting DDR1^17^. In addition, exogenous miR-199b-5p inhibited the growth of bone marrow mesenchymal stem cells (MSCs) and promoted the differentiation of bone MSCs into chondrocytes by targeting the *JAG1* pathway ^18;19^. Our functional experiments showed that overexpression of miR-199b-5p reduced chondrocyte viability and the expression of anabolic factors such as COL2A1, AGGREGN, and SOX9. It also increased inflammation levels and decreased pain thresholds in control mice. Although no pathological changes were observed in the articular cartilage of control mice, a similar study demonstrated that pathological cartilage changes only occurred after six months in mice with miR-211 and miR-204 knockout^20^. Therefore, we speculate that the time of overexpression of miR-199b-5p in our experiment was too short, so it only caused inflammation and pain threshold response. To best of our knowledge, this is the initial investigation documenting the involvement of miR-199b-5p in KOA.

*Fzd6* is known to be up-regulated during the osteogenic differentiation of MSCs and can be regulated by miR-194-5p to activate the WNT signaling pathway, thereby promoting osteogenic differentiation of MSCs^21^. *Gcnt2* has been found to induce epithelial-mesenchymal transition and enhance migration and invasion of esophageal squamous cell carcinoma cells^22^. Our findings indicate that miR-199b-5p plays a crucial role in KOA by potential targeting *Fzd6* and *Gcnt2*. However, whether miR-199b-5p truly functions through *Fzd6* and/or *Gcnt2* requires genetic knockdown of *Fzd6* and *Gcnt2* in the presence of miR-199b-5p. Future research should further explore the roles and mechanisms of *Fzd6* and *Gcnt2* in the context of KOA.

This study also has some limitations. We initially detected serum exosomes miRNA and later examined the miRNA through *in vitro* experiment and animal study. However, whether the miRNAs targeting the joints have the same role as the serum exosome miRNAs or blood miRNA has not been clear. We also performed direct intra-articular injection of adenoviral vectors. The intra-articular injection can reduce the exposure of extra-articular tissue, thereby minimizing the associated side effects and improving targeting. In this study, the chondrocyte experiments were conducted in a 2D manner, which may lead to chondrocyte de-differentiation and thus may weaken the conclusion of the chondrocyte response to the treatments. Therefore, in the future study, we will adopt 3D culture system for experiments. Besides, more in vivo study such as gene knockout mice study can be used in the future study.

In conclusion, we found that miR-199b-5p is elevated in osteoarthritis and may affect cell viability and related cytokines by potential targeting *Gcnt2* and *Fzd6*. Overexpression of miR-199b-5p induced OA-like pathological changes in normal mice and inhibiting miR-199b-5p alleviated symptoms in KOA mice, suggesting it may be a target for treatment of OA. Through human clinical trials, cell experiments, and animal models, we not only identified a new OA-related miR-199b-5p but also examined the biological function of miR-199b-5p in OA.

## MATERIALS AND METHODS

### Human samples

Study participants were recruited from the Hospital of Chengdu University of Traditional Chinese Medicine and the surrounding communities. The study was approved by the Ethics Review Committee of the Hospital at Chengdu University of Traditional Chinese Medicine (2016KL-017) and conformed to the ethical guidelines of the 1975 Declaration of Helsinki. Serum was collected from all participants and serum samples were stored at -80°C.

### Human subjects

Patients were enrolled if they fulfilled the following criteria: (1) diagnosed with KOA according to the ACR; (2) age between 40 and 65 years; (3) agreed to cooperate with researchers in all research procedures after enrollment; (4) provided with written informed consent. Patients with any of the following conditions were excluded: (1) accompanied with other diseases such as rheumatoid arthritis, bone tumors, and bone tuberculosis; (2) treated with intra-articular gluco-corticoid or viscoelastic supplementation in the last six months; (3) knee replacement history; (4) had complicated cardiovascular disease, diabetes, skin disease, and liver or kidney impairment; (5) pregnant or breastfeeding women; (6) accompanied with mental and intellectual disabilities; or (7) undergoing other clinical trials.

### Extraction and sequencing of serum exosome miRNAs

Serum samples were filtered using 0.22 µM filters. Exosomes were isolated from the serum sample using ExoQuick™ Exosome Precipitation Solution (System Biosciences) following the manufacturer’s instructions. Briefly, serum was thawed on ice and centrifuged at 4000 rpm for 15 min to remove any cells or cellular debris. Next, 50 μL ExoQuick Solution was added into the 200 μL serum sample and mixed thoroughly. The exosomes were suspended in PBS.

Exosomes were characterized by electron microscopy (Tecnai G2 Spirit 120KV, FEI), nanoparticle tracking analysis (NTA) (NTA 3.2 Dev Build 3.2.16), and western blot analysis (for CD9, CD63 and CD81). Sequencing was performed by single-end sequencing (1×150 bp) on Illumina NextSeq 500. The libraries were sequenced on an Agilent 2100 Bioanalyzer platform. The mirdeep2 software (https://www.mdc-berlin.de/content/mirdeep2-documentation) was used to analyze the miRNA sequences and quantification. The heatmap was plotted based on the log2 (fold change), using Heatmap Illustrator software (Heml 1.0).

### Total RNA was isolated from human serum, cell and mice

Chondrocytes using a kit (Yeasen,Shanghai, China) according to the manufacturer’s instructions. Reverse transcription was performed using 1000 ng total RNA and a Prime Script RT Reagent Kit(Yeasen, Shanghai, China) or Prime Script RT Master Mix (Yeasen, Shanghai, China), which were used for miRNA and mRNA, respectively. For miRNA, the reactions were incubated at 42°C for 15 min followed by inactivation at 85°C for 5 s. qRT-PCR amplification was assessed in a CFX Connection Real-Time System (Bio-Rad) using the SYBR Premix Ex Taq II kit (Yeasen, Shanghai,China). The following cycling conditions were used: 95°C for 30 s, followed by 40 cycles of 95°C for 5 s and 60°C for 30 s. All reactions were performed in duplicate and normalized to the internal reference U6 for miRNA and GAPDH mRNA for mRNAs. The 2^-ΔΔCT^ CT method was used to evaluate the relative mRNA/miRNA expression levels.

### RNA primer

Primers are listed in Supplementary Table 1.

### Bioinformatics analysis

GO analysis was performed, involving the biological process (BP), cellular component (CC), and molecular function (MF), using DAVID^23^ (https://david.ncifcrf.gov/home.jsp).We used the miRNA target gene prediction websites miRDB^24^(http://mirdb.org/), miRWalk^25^(http://mirwalk.umm.uni-heidelberg.de/), Starbase^26^ (https://starbase. sysu.edu.cn/), DIANA-TarBase^27^ (http://diana.imis.athena-innovation.gr/DianaTools/ index.php?r=tarbase/index), Targetscan^28^ (https://www.targetscan.org/vert_72/). The target genes of mmu-miR-199b-5p were predicted, and the intersection of the target gene prediction list results of the five sites was taken.

The dysregulated miRNAs were compared to relevant published miRNA data from human KOA patients. The dataset GSE105027 was obtained from serum samples of KOA patients, while GSE175961 comprised sequencing data from KOA patient cartilage. Finally, we validated the expression of the target genes Gcnt2 and Fzd6 in KOA patients using synovial data (GDS5401, GDS5403) and cartilage data (GDS3758). All operations will be performed using the R software.

### Primary mouse chondrocyte culture and other cell experiments

According to the protocol by Gosset et al^29^. young mice 3-5 days after birth were taken, cartilage tissues were extracted, and the cells were cultured in complete medium after digestion with collagenase. The cells were cultured in F12 medium containing 10% fetal bovine serum and 1% penicillin-streptomycin at 37°C and a 5% CO_2_. The medium was changed every three days, and cells were cultured to the second passage for experiments. Throughout the experiment, we used second-generation chondrocytes and measured cell viability using the CCK-8 assay. The stimulation time for lipopolysaccharide-induced inflammatory damage model was 6 hours. Regarding virus infection, the cells were infected after 36h of passage when their density reached approximately 30%. According to the virus operation manual, the total infection time was approximately 40 h.

### Animal model of KOA

Animal experiments were performed using total of 108 8-week-old male C57BL/6 mice. The protocol was approved by the Committee on the Ethics of Animal Experiments of Chengdu University of Traditional Chinese medicine (2019-04). After randomization, the KOA mouse model was established by orthotopic injection of MIA (sigma) into the knee joint of mice. After the animals were anesthetized with isoflurane in a small animal anesthesia machine, the right knee joint of the mouse was shaved, the knee joint was flexed 90°, and 10 μL of MIA solution was injected into the knee joint cavity using a 28G microsyringe. The syringe needle was slowly withdrawn and the knee joint was gently moved. Baseline testing of behavioral pain thresholds was conducted prior to model establishment. After model established, the behavioral pain threshold testing was on days 3, 7, 10, 14, 21, and 28 after model establishment. After the 3rd day and 14th day after the start of the experiment, the experimental mice were injected with adenovirus (total volume 10μL) in the knee joint, and the control mice were injected with empty adenovirus as a control (Hanbio tech, Shanghai, China)^30^.

### Thermal pain threshold detection

A pain threshold detection instrument was used to measure the latency of the right plantar leg raising reflex of mice under heat radiation. Three days before the experiment, mice were cut off from water for half a day and adapted to the environment for 1 h. A certain intensity of pyrogen was used to irradiate the plantar position of the mouse’s right limb, and a machine was used to automatically record the time when the mouse moved the limb from thermal pain. The resting intensity of the thermal pain stimulator was set to 10%, and the maximum duration was set to 20 s to avoid prolonged heating and burning of skin. After the light source is aimed at the sole of the mouse, we observed the mouse in real time. If paw raising, paw licking, or paw retraction was observed, the irradiation was stopped and the irradiation time of the instrument was recorded. The right limb of each mouse was tested five times, with an interval of 10 min each time; outliers were eliminated, and the average value was calculated and included in the statistics^31^.

### ELISA and μCT scanning

Mice were sacrificed after four weeks, and serum was collected for ELISA assay (Jiangsu Jingmei Biotechnology Co., Ltd., Yancheng, China). The knee joint specimens of the right hindlimb of mice were scanned using a high-resolution micro-CT skysan1267 instrument (Bruker, Germany). The sample was removed from fixative and dried. The scanning parameters were as follows: voltage 55 KV, current 200 μA, and filter 0.25AL. After scanning, three-dimensional reconstruction was performed using NRecon1.7.4.2 software.

### Safranin Fast green staining

Mouse knee joint staining was performed using the Safranin Fast Green Staining Kit (Servicebio Biological Technology, Wuhan, China). The sample was examined under a microscope, and the cartilage integrity was scored according to the OARSI grading system, in which the score ranged from 0 to 6 points^32^.

### Luciferase assay

We utilized Targetscan to predict potential binding sites and designed sequences accordingly. The wild-type (WT) and mutant-type (MUT) sequences (according to the predicted binding site) were inserted into the pmiRGLO plasmid. HEK-293T cells were seeded in 6-well plates at 24 h before transfection. GCNT2 and FZD6 3′UTR-wt and gcnt2 and fzd6 3′UTR-mut plasmids (500 ng) and 20 nmol miR-199b-3p and NC were co-transfected with Lipofectamine 3000 (Hanbio Biotechnology, China) following the manufacturer’s instructions. After 48h, firefly and Renilla luciferase activities were calculated using the Promega Dual-Luciferase system following the manufacturer’s instructions (Hanbio tech, Shanghai, China). Firefly/Renilla luciferase was measured to evaluate relative luciferase activity.

### Comparative analysis of sequence conservatism between human and mouse

Firstly, the sequence information of mmu_miRNA-199b-5p was used to locate the human homologous sequence in the UCSC^33^ database. Based on this positional information and the source gene, a further comparison was conducted in miRbase^34^ to identify the nearest miRNA at the position of the human genome.

### Statistical analysis

Data are reported as the means ± SD. Normal distribution and homogeneity of variance of data were first tested. Shapiro-Wilk test was used to verify data normality, while Levene’s test or Browne-Forsythe test was adopted for assessment of variance equality. Statistical analysis was performed by unpaired two *t*-test and Tukey-corrected one-way analysis of variance (ANOVA) for comparisons between groups. Tukey-corrected two-way ANOVA for the comparison of mice thermal pain threshold data. *P* < 0.05 was considered statistically significant for all statistical calculations. PRISM 8.0 (GraphPad Software, San Diego, CA, USA) was used for data analysis.

## Acknowledgements

This work was supported by the National Key R&D Program of China (No. 2022YFC3500703); Fund of Science and Technology Department of Sichuan Province, China (No.2022ZDZX0033); and Xinglin Scholar Foundation of Chengdu University of Traditional Chinese Medicine (No. QNTD2022003).

## Contributions

F.T., Z.Q., and Q.-F.W. designed the study.

F.T. and Y.-L.P. performed most in vitro and in vivo experiments.

F.T. and J.-M.W. analyzed data.

Z.Q., M.-Q.T. and S.-H.L. performed the miRNA sequencing and interpreted the data.

Z.Q. collected and inspected human patient samples.

F.T. and Q.-F.W. drafted the manuscript.

All authors read and edited the manuscript.

F.-R.L., S.-G.Y., X.W, S.-H.Z. and Q.-F.W. supervised the study.

## Competing interests

The authors declare that they have no known competing financial interests or personal relationships that could have appeared to influence the work reported in this paper

## Data Sharing Statement

All data are available in the main text or the supplementary materials.

## List of Supplementary Materials

Present a list of the Supplementary Materials in the following format. Materials and Methods

Fig S1. Identification of exosome. (A) Transmission electron microscope scanning of isolated exosomes from the serum of participants. (B) The Nano sight particle analysis of isolated exosomes from the serum of participants. (C) The Western blot analysis of symbolic surface markers of isolated exosomes from the serum of participants.

Fig S2. Fluorescence of Adenovirus-Infected Chondrocytes. Fig S3. Expression of adenovirus in mouse knee joint.

Fig S4. Establishment of KOA model in mice by injection of MIA. (A) Behavioral detection of animal thermal pain threshold. (B) Microscopic observation of the surface of the mouse knee joint. (C) Safranin-fast green staining and semiquantitative scoring of articular cartilage. (D) 3D reconstruction images of joints from μCT scans. \**P*<0.05, \*\**P*<0.01, *n*=6.

Fig S5. Venn diagram showing differentially expressed miRNAs in the OA group compared with healthy patients and patients who recovered after acupuncture treatment.

Fig S6. Comparative analysis of sequence conservatism between human and mouse. (A) By using the sequence information of mmu_miRNA-199b-5p, we located the position of its human homologous sequence in the UCSC database. (B) Based on the positional information and the source gene, we further aligned this position with the closest miRNA in miRbase. (C) We compared the sequences of hsa_miR-199b-5p and mmu_miR-199b-5p.(D) Conservation analysis was performed to compare the sequence conservation of miR-199b-5p.

Fig S7. Primary mice chondrocytes we cultured (P1) and the secondary generation cells (P2) we used in the following experiment.

Supplement file8. We predicted the potential binding sites of miRNA-199b-5p in the 3’-untranslated regions (UTRs) of two target genes, Fzd6 and Gcnt2, in both human and mouse.

**Supplement table 1:**
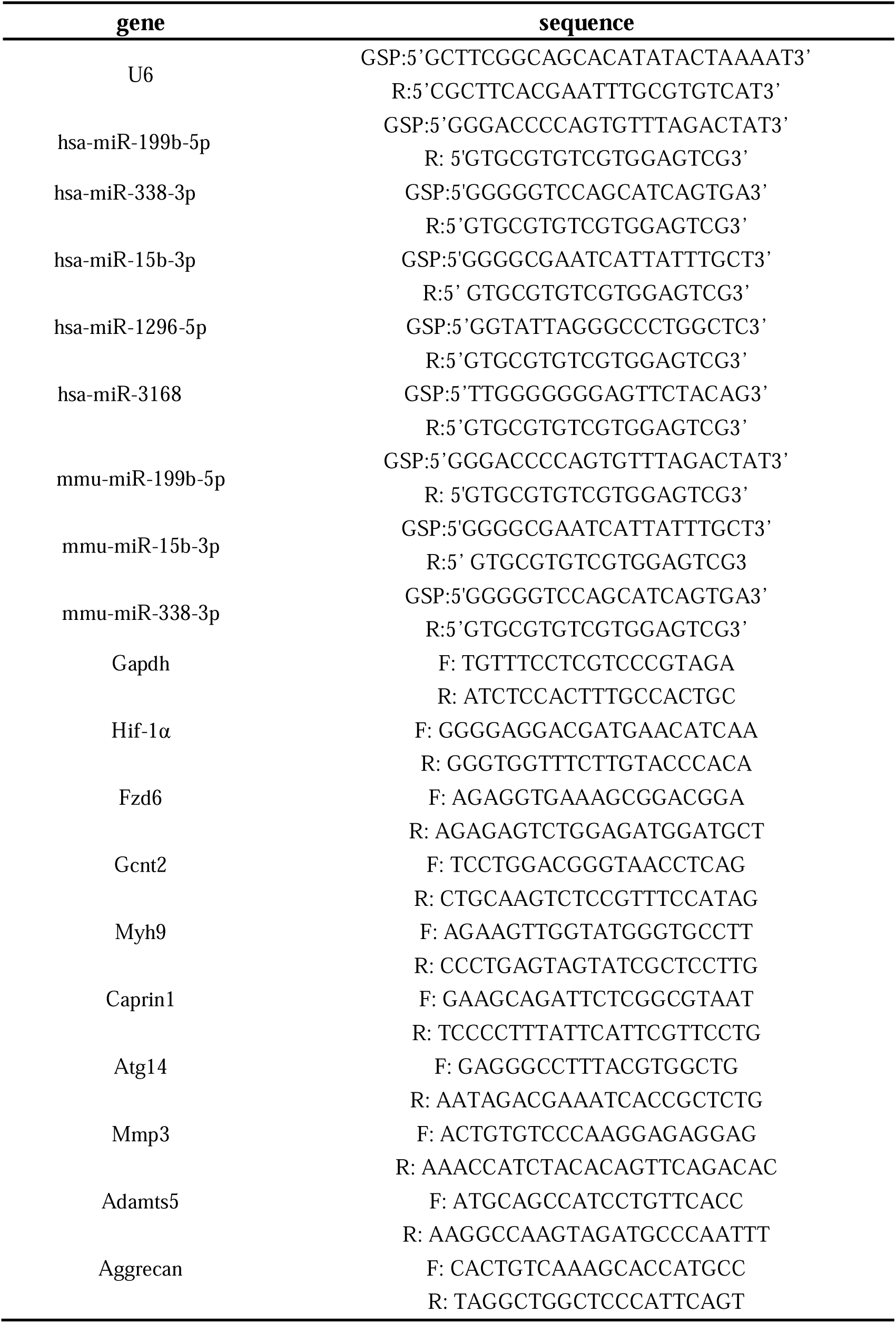
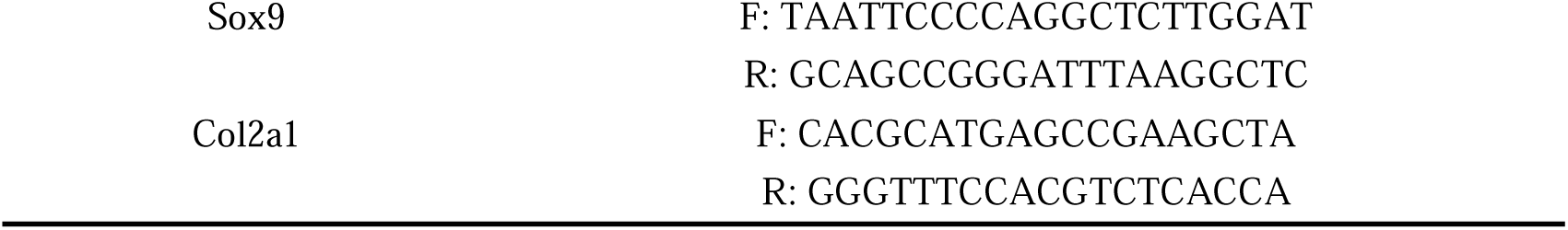
Primer List.

**Supplement table 2:** Basic information for recruiting patients.

| Baseline | KOA ( N=15 ) | HC ( N=10 ) | *P* value |
| --- | --- | --- | --- |
| Age | 52.67 ± 7.35 | 51.50 ± 7.71 | 0.706 |
| Height | 1.63±0.07 | 1.60±0.05 | 0.201 |
| Weight | 65.03±9.53 | 61.30±8.47 | 0.327 |
| BMI | 24.28 ± 2.39 | 23.90± 2.65 | 0.715 |

---

**Dear editor:**

We appreciate very much for the opportunity to reconsider our paper to be published in the *eLife*.

We thank the constructive criticism provided by the reviewers and editor. Based on these suggestions, we have thoroughly reworked the manuscript. More specifically but not limit:

1) We have corrected the mistakes mentioned by the reviewers on a point-by-point basis.

2) We have provided additional experimental evidences to explain the rationale behind selecting five miRNAs for q-PCR validation. Furthermore, we have elaborated on the reasons for focusing primarily on research related to cartilage.

3) In response to concerns regarding overinterpretation in the manuscript, we have made more precise descriptions and revisions. Furthermore, we have added some details in our methods, including the addition of results showing the conservation of miR-199b-5p sequences between human and mouse species.

4) We have provided additional details on the experiments, including the process for predicting target genes, timing of chondrocyte culture and other experimental operations.

5) Finally, we have made additional revisions to the details of the figures to avoid any distortions and enhance the precision of the language.

Below please find our responses to the reviewers’ comments on a point-by-point basis. You also can track the changes in the modified manuscript. We believe that this revision has been substantially improved. We do hope that our revised manuscript merits to be published in the *eLife*.

Please do not hesitate to contact us if you require any additional information. Best regards,

Qiao-feng Wu,

M.D, Professor,

Acupuncture and Moxibustion College, Chengdu University of Traditional Chinese Medicine

*eLife assessment*

The manuscript provides interesting evidence that miR-199b-5p regulates osteoarthritis and as such it may be considered as a potential therapeutic target. This finding may be useful to further advance the field.

***<u>Response:</u>***Thank you for your positive comments.

Although the study is considered potentially clinically relevant, the evidence provided was deemed insufficient and incomplete to support the conclusions drawn by the authors.

***<u>Response:</u>***Thank you for your critical comments and constructive advices. We have response point to point according to the reviewers’ questions and thoroughly re-working our manuscript. We hope the revised manuscript can be qualified to the criteria and be published on the journal of *eLife*.

**Reviewer #1 (Public Review):**

Summary:

In this manuscript, the authors observed that miR-199b-5p is elevated in osteoarthritis (OA) patients. They also found that overexpression of miR-199b-5p induced OA-like pathological changes in normal mice and inhibiting miR-199b-5p alleviated symptoms in knee OA mice. They concluded that miR-199b-5p is not only a potential micro-target for knee OA but also provides a potential strategy for the future identification of new molecular drugs.

***<u>Response:</u>***Thanks for your comment.

Strengths:

The data are generated from both human patients and animal models.

***<u>Response:</u>***Thanks for the positive comment.

Weaknesses:

The data presented in this manuscript is not solid enough to support their conclusions. There are several questions that need to be addressed to improve the quality of this study.

The following questions that need to be addressed to improve the quality of the study.

1. Exosomes were characterized by electron microscopy and western blot analysis (for CD9, 264 CD63, and CD81). However, figure S1 only showed two sample WB results and there is no positive and negative control as well as the confused not clear WB figure.

***<u>Response:</u>***Thank you for your suggestion. We acknowledge that a comprehensive identification of extracellular vesicles should include both positive and negative samples. However, in some of the initial studies we referenced, the positive and negative control were not mentioned^1;2^. In our study, we identified extracellular vesicles using a combination of electron microscopy, nanoparticle tracking analysis, and marker detection of exosomes. We agree that having negative samples would make our results more convincing, and we will include a negative control group in our future experiments. Additionally, we have provided clearer images in the revised version. (supplemental fig1 A)

**Reference**

1) Ying W, Riopel M, Bandyopadhyay G, et al. Adipose Tissue Macrophage-Derived Exosomal miRNAs Can Modulate In Vivo and In Vitro Insulin Sensitivity. Cell. 2017;171(2).

2) Fang T, Lv H, Lv G, et al. Tumor-derived exosomal miR-1247-3p induces cancer-associated fibroblast activation to foster lung metastasis of liver cancer. Nature Communications. 2018;9(1):191.

2. The sequencing of miRNAs in serum exosomes showed that 88 miRNAs were upregulated and 89 miRNAs were downregulated in KOA patients compared with the control group based on fold change > 1.5 and p < 0.05. Figure 2 legend did not clearly elucidate what those represent and why the authors chose those five miRNAs to further validate although they did mention it with several words in line 108 ‘based on the p-value and exosomal’.

***<u>Response :</u>*** In fact, our study included two additional groups: the acupuncture treatment group (4 weeks of continuous acupuncture treatment) and the waiting treatment group (no intervention, followed by acupuncture treatment after 4 weeks), in addition to the healthy control and knee osteoarthritis (OA) patient groups. After comparing these four groups, we found that 11 genes (hsa-miR-504-3p, hsa-miR-1915-3p, hsa-miR-103a-2-5p, hsa-miR-887-3p, hsa-miR-1228-5p, hsa-miR-34c-3p, hsa-miR-3168, hsa-miR-518e-3p, hsa-miR-1296-5p, hsa-miR-338-3p, and hsa-miR-199b-5p) were upregulated in KOA patients but downregulated after acupuncture treatment, with no change in the waiting treatment group. Additionally, 7 genes (hsa-miR-448, hsa-miR-514a-3p, hsa-miR-4440, hsa-let-7f-5p, hsa-let-7a-5p, hsa-let-7d-5p, and hsa-miR-15b-3p) were downregulated in KOA patients but upregulated after acupuncture treatment, with no change in the waiting treatment group. Considering the improvement in clinical symptoms of KOA patients after acupuncture treatment, we believe that these 18 genes are of significant value. Based on overall expression abundance and species specificity, we finally selected 5 genes, namely the 5 genes mentioned in this article. Regarding this result, we have already included it in the supplementary fig5(fig. S5).

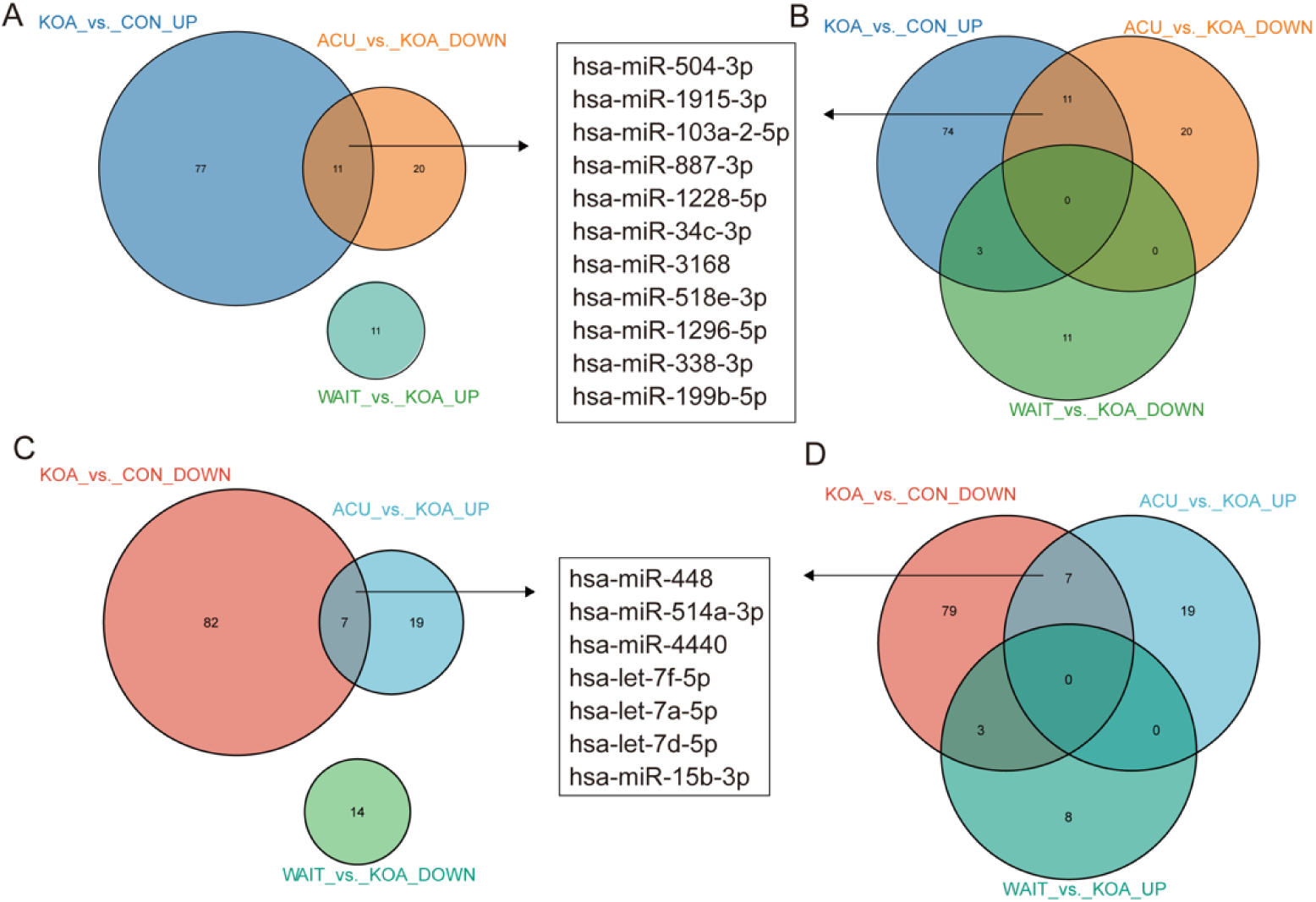
Venn diagram showing differentially expressed miRNAs in the OA group compared with healthy patients and patients who recovered after acupuncture treatment.

3. In Figure 3 legend and methods, the authors did not mention how they performed the cell viability assay. What cell had been used? How long were they treated and all the details? Other figure legends have the same problem without detailed information.

***<u>Response:</u>***Thank you for your suggestions. In Figure 3, cell viability was determined using the CCK-8 assay. We used second-generation chondrocytes for this analysis. The chondrocytes were obtained from young mice aged 3-5 days after birth. The cartilage tissues were extracted, and the cells were cultured in complete medium after digestion with collagenase. The detailed description of the cell viability assay, cell culture procedures, specific timing, and treatment methods of the cells used can be found in our revised manuscript. (page14-15, line304-313)

Besides, we have made thorough revisions to all figure legends to provide a clearer explanation of the relevant content.

4. The authors claimed that Gcnt2 and Fzd6 are two target genes of miR-199b-5p. However, there is no convincing evidence such as western blot to support their bioinformatics prediction.

***<u>Response:</u>***In the current study, we first identified six potential target genes by intersecting the predicted targets obtained from six bioinformatics websites. Subsequently, q-PCR was employed to test all six genes, revealing two genes with significant changes, namely *Fzd6* and *Gcnt2*. We then predicted the binding sites of these genes and validated their existence through luciferase assays. Moreover, we examined the expression of these two potential targets in human KOA samples using a human database and found them to be expressed specifically in the samples. These results suggest that *Fzd6* and *Gcnt2* are potential target genes for KOA. However, we didn’t do western blot assay to verify the results. Based on your suggestions, we have further discussed the limitations of our study in this regard and proposed future research strategies.

5. To verify the binding site on 3’UTR of two potential targets, the authors designed a mouse sequence for luciferase assay, but not sure if it is the same when using a human sequence.

***<u>Response:</u>***Thank for your great advice. We carried out the comparative analysis of sequence conservatism between human and mouse, and find the binding site on 3’UTR matches to human sequence very well. The sequence conservation between hsa_miR-199b-5p and mmu_miR-199b-5p was as high as 95.65%. We added the methods and results in the revised manuscript. (page9, line181-184; page17, line361-365) (supplemental fig6)

In detail: Firstly, the sequence information of mmu_miRNA-199b-5p was used to locate the human homologous sequence in the UCSC database. The homologous sequence was found to be located in the human genome at chr9:128244721-128244830 (supplemental fig6 A). Based on this positional information and the source gene, a further comparison was conducted in miRbase to identify the nearest miRNA at the position of the human genome. It was discovered that hsa_miR-199b-5p is positionally conserved and located at chr9:128244721-128244830 (supplemental fig6 B). The sequence of hsa_miR-199b-5p was obtained from the miRbase database (supplemental fig6 C), and a comparative analysis was performed between the sequences of humans and mouse (supplemental fig6 D). Besides being positionally conserved, the sequence conservation between hsa_miR-199b-5p and mmu_miR-199b-5p was as high as 95.65%, indicating a good sequence conservation.

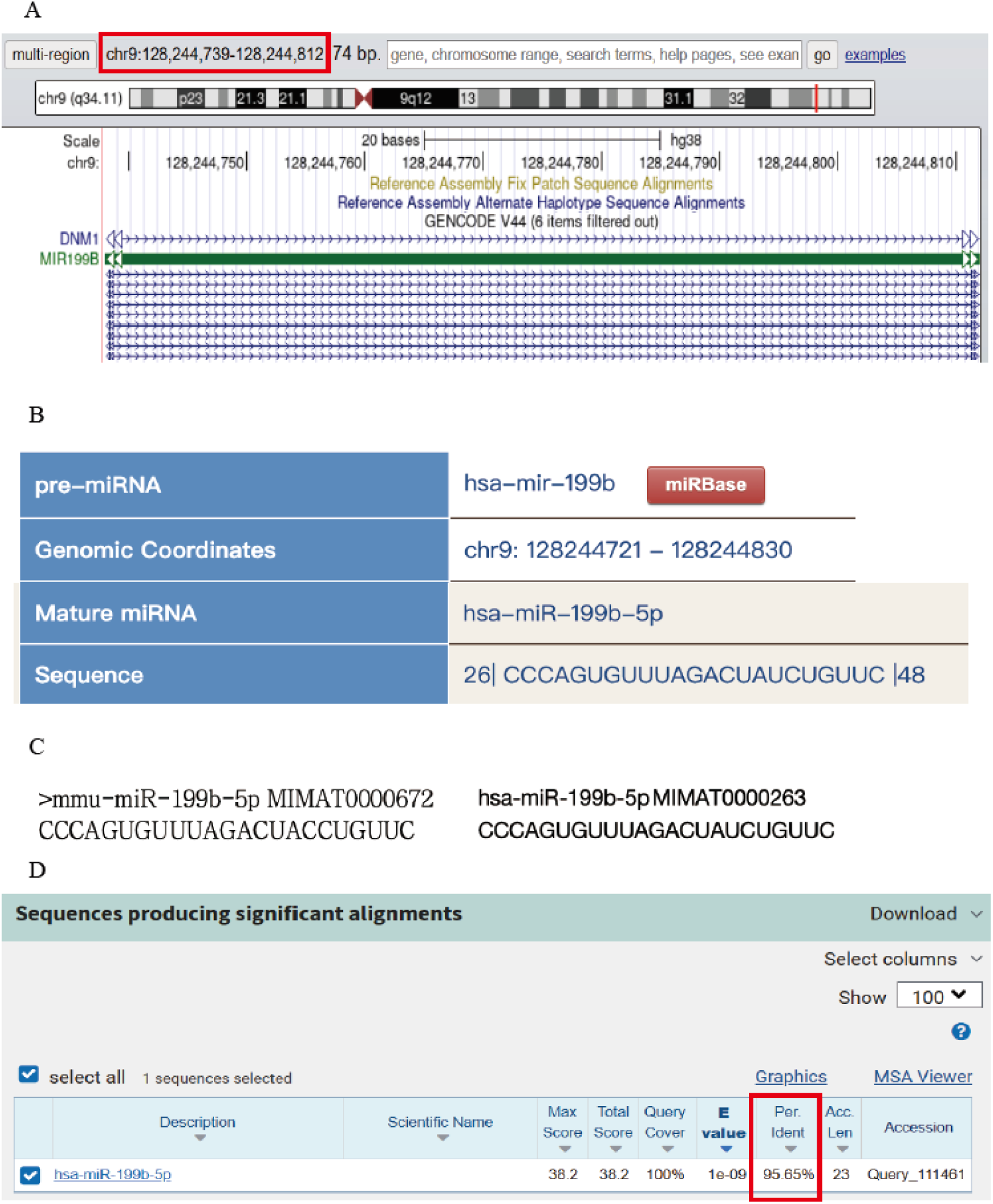
(A) By using the sequence information of mmu_miRNA-199b-5p, we located the position of its human homologous sequence in the UCSC database. (B) Based on the positional information and the source gene, we further aligned this position with the closest miRNA in miRbase. (C) We compared the sequences of hsa_miR-199b-5p and mmu_miR-199b-5p. (D) Conservation analysis was performed to compare the sequence conservation of miR-199b-5p.

**Reviewer #2 (Public Review):**

Summary:

The authors identified miR-199b-5p as a potential OA target gene using serum exosomal small RNA-seq from human healthy and OA patients. Their RNA-seq results were further compared with publicly available datasets to validate their finding of miR-199b-5p. In vitro chondrocyte culture with miR-199b-5p mimic/inhibitor and in vivo animal models were used to evaluate the function of miR-199b-5p in OA. The possible genes that were potentially regulated by miR-199b-5p were also predicted (i.e., *Fzd6* and *Gcnt2*) and then validated by using Luciferase assays.

***<u>Respons:</u>***We greatly appreciate Reviewer #2 constructive comments. Strengths:

1. Strong in vivo animal models including pain tests.

2. Validates the binding of miR-199b-5p with *Fzd6* and binding of miR-199b-5p with *Gcnt2*.

***<u>Response:</u>***Thanks for positive comment.

Weaknesses:

1. The authors may overinterpret their results. The current work shows the possible bindings between miR-199b-5p and *Fzd6* as well as bindings between miR-199b-5p and *Gcnt2*. However, whether miR-199b-5p truly functions through *Fzd6* and/or *Gcnt2* requires genetic knockdown of *Fzd6* and *Gcnt2* in the presence of miR-199b-5p.

***<u>Response:</u>***In this study, we employed a comprehensive approach by integrating data from six bioinformatics databases to identify potential target genes for miR-199b-5p. Subsequent qPCR analysis revealed significant changes in two genes, *Fzd6* and *Gcnt2*. We then utilized luciferase assays to validate the predicted binding sites and confirmed the interaction between miR-199b-5p and these genes. Additionally, we examined the expression profiles of these potential target genes in human KOA samples using a human database, which unveiled distinct expression patterns.

While our findings suggest that *Fzd6* and *Gcnt2* may serve as potential target genes for miR-199b-5p, we acknowledge the necessity for further experimental validation and in-depth functional characterization. Building upon your insightful recommendations, we have thoroughly addressed the research limitations and proposed potential research strategies for future investigations in our discussion.

(page11, line227-231)

2. In vitro chondrocyte experiments were conducted in a 2D manner, which led to chondrocyte de-differentiation and thus may not represent the chondrocyte response to the treatments.

Response:we admit that 3D culture system will be more accurate and reliable.

However, according to Liu Qianqian et al researches^3^, the 2D culture systems were also used and work well. Besides, the second-generation primary mice chondrocytes we used in the current study did not exhibit a significant dedifferentiated morphology. So, considering the experiment condition in our lab, we chose the second-generation cultured primary mouse chondrocytes in the whole process of cell experiment. To show the reliability of the cells, we provided more pictures in the supplement fig 7(fig. S7) In the future study, we will adopt 3D culture system for experiments. Thank you for your advices and we have added this limitation in the revised manuscript. (page11, line237-240)

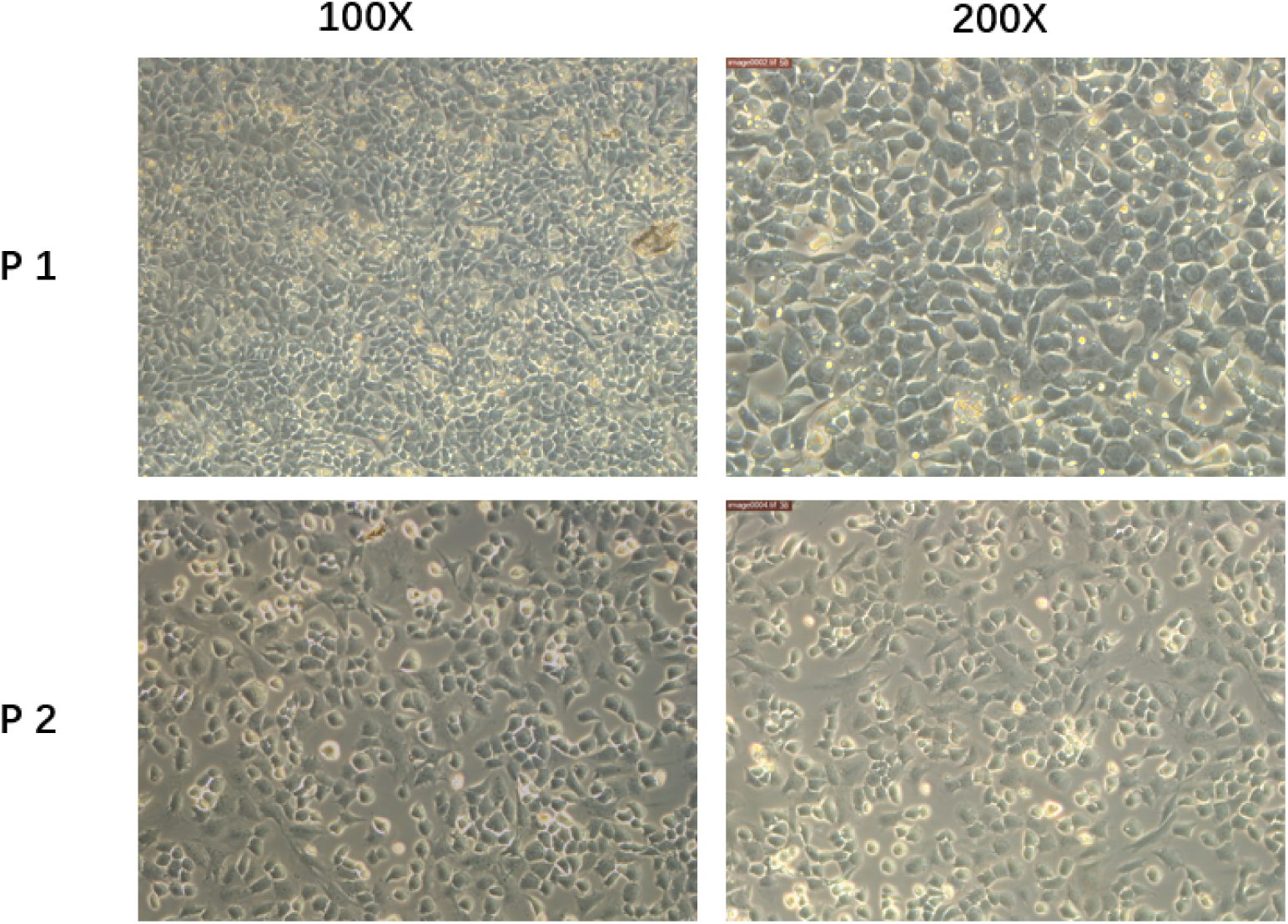
Primary mice chondrocytes we cultured (P1)and the secondary generation cells(P2) we used in the following experiment.

**References which used 2D** :

3. Liu Q, Zhai L, Han M, et al. SH2 Domain-Containing Phosphatase 2 Inhibition Attenuates Osteoarthritis by Maintaining Homeostasis of Cartilage Metabolism via the Docking Protein 1/Uridine Phosphorylase 1/Uridine Cascade. Arthritis & Rheumatology (Hoboken, NJ). 2022;74(3):462-474.

3. There is a lack of description for bioinformatic analysis.

***<u>Response:</u>***Sorry for our neglection. We have added relevant descriptions and details. (Pages 14, line299-303)

4. There are several errors in figure labeling.

***<u>Response:</u>***We have revised. (Fig. 3, Fig. 4, Fig. 5 and Fig. 7)

Recommendations for the authors: please note that you control which revisions to undertake from the public reviews and recommendations for the authors

***<u>Response:</u>***We appreciate the reviewers’ feedback as we believe it has significantly contributed to the refinement of our manuscript. We are confident that our revisions have strengthened the quality and impact of our study, and we agree that the suggestions presented by the reviewers are valuable and appropriate for publication.

**Reviewer #2 (Recommendations For The Authors):**

I would like to thank the authors for investigating the functional role of miR-199b-5p in knee OA. While this study has the potential to provide valuable knowledge to the fields of miRNAs and joint diseases, significant improvements in several areas are required.

***<u>Response :</u>*** We appreciate your constructive comments, and we have made a substantial improvement to the manuscript. We thank all the reviewers for their advice as well as their criticisms.

Major concerns:

1. According to the Authors, miR-199b-5p is identified by the results from their own miRNA-sequencing as well as comparison with other publicly available datasets (both synovium and cartilage datasets). It is unclear to me why the synovium dataset was used here as it appears that the entire manuscript was mainly focused on chondrocytes.

***<u>Response:</u>***Thank you for your question. As we are aware, cartilage degradation is the initial pathological change in knee osteoarthritis (KOA), which subsequently leads to other pathological changes such as synovial inflammation^4^. These factors are interrelated, and current research on KOA encompasses cartilage, synovium, and system inflammation et al. Therefore, when we identified a large number of dysregulated miRNAs in extracellular vesicles isolated from serum, it was crucial to determine whether these dysregulated miRNAs were also altered in cartilage or synovium. To address this, we compared our findings with publicly available databases and found a higher overlap with the cartilage cell dataset, including miRNA-199b. Consequently, we decided to focus our subsequent investigations on cartilage-related research.

**Reference**

4. Hunter D, Bierma-Zeinstra S. Osteoarthritis. Lancet (London, England). 2019;393(10182):1745-1759.

2. Also, 169 of 177 differentially expressed exosome miRNAs were intersected with differentially expressed miRNAs from OA cartilage datasets. It is surprising that in the 5 selected miRNAs for further qRT-PCR validation, 3 out of 5 were not in the exosome miRNA dataset (i.e., hsa-mir-1296-5p, hsa-mir-15b-3p, and hsa-mir-338-3p; page 5, line 109 and Fig. 1B). Isn’t that selecting the miRNAs that both differently expressed in exosome and cartilage datasets for validation more essential? Furthermore, from the Authors’ exosome miRNA dataset, only 5 out of 15 KOA patients actually exhibited up-regulated miR-199b-5p vs. health controls. Please elaborate on how the target was determined.

***<u>Response :</u>*** In fact, our study included two additional groups: the acupuncture treatment group (4 weeks of continuous acupuncture treatment) and the waiting treatment group (no intervention, followed by acupuncture treatment after 4 weeks), in addition to the healthy control and knee osteoarthritis (OA) patient groups. After comparing these four groups, we found that 11 genes (hsa-miR-504-3p, hsa-miR-1915-3p, hsa-miR-103a-2-5p, hsa-miR-887-3p, hsa-miR-1228-5p, hsa-miR-34c-3p, hsa-miR-3168, hsa-miR-518e-3p, hsa-miR-1296-5p, hsa-miR-338-3p, and hsa-miR-199b-5p) were upregulated in KOA patients but downregulated after acupuncture treatment, with no change in the waiting treatment group. Additionally, 7 genes (hsa-miR-448, hsa-miR-514a-3p, hsa-miR-4440, hsa-let-7f-5p, hsa-let-7a-5p, hsa-let-7d-5p, and hsa-miR-15b-3p) were downregulated in KOA patients but upregulated after acupuncture treatment, with no change in the waiting treatment group. Considering the improvement in clinical symptoms of KOA patients after acupuncture treatment, we believe that these 18 genes are of significant value. Based on overall expression abundance and species specificity, we finally selected 5 genes, namely the 5 genes mentioned in this article. Regarding this result, we have already included it in the supplementary fig5(fig. S5).

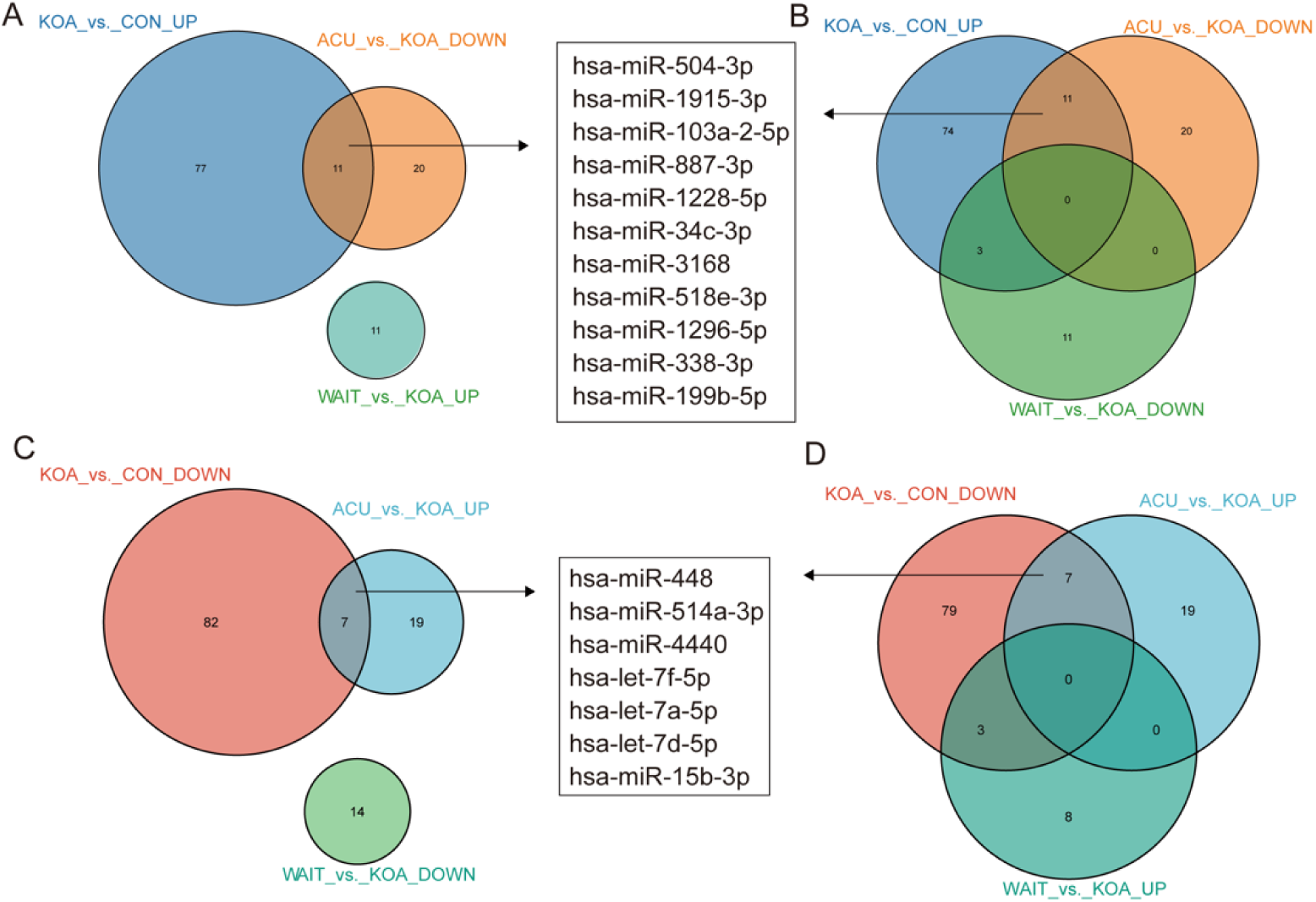
Venn diagram showing differentially expressed miRNAs in the OA group compared with healthy patients and patients who recovered after acupuncture treatment.

3. There is also a lack of description for bioinformatic analysis regarding how miRNA sequencing datasets were analyzed. What R/python packages or algorithms were used? What were the QC criteria?

***<u>Response:</u>***We apologize for any confusion caused. We have now included a clear description of the method employed, and R was utilized for this data analysis (revised in Page14, Line301-305). To ensure consistency, we compared our findings with publicly available human serum data from the database (GSE105027) using a fold change threshold of > 1.5 and a significance level of *p* < 0.05. In the cartilage data (GSE175961), we observed a list of miRNAs with shared expression patterns, yet the precise differential values could not be determined.

4. Another major concern is the chondrocyte culture method. Chondrocytes should be cultured in a 3D manner (i.e., a 3D pellet culture system or a micro mass culture method). 2D cultured chondrocytes tend to de-differentiate into MSC-like cells and thus lose their chondrocyte phenotype. This is evident from Fig. 3B and C. Cells started to spread out and only a few cells were positive for COL2A1 with a deep brown staining color. Thus, the results from the *in vitro* studies may not be representative of chondrocyte response to the treatments.

***<u>Response:</u>***we admit that 3D culture system will be more accurate and reliable. However, according to Liu Qianqian et al researches^3^, the 2D culture systems were also used and work well. Besides, the second-generation primary mice chondrocytes we used in the current study did not exhibit a significant dedifferentiated morphology. So, considering the experiment condition in our lab, we chose the second-generation cultured primary mouse chondrocytes in the whole process of cell experiment. To show the reliability of the cells, we provided more pictures in the supplement fig 7(fig. S7) In the future study, we will adopt 3D culture system for experiments. Thank you for your advices and we have added this limitation in the revised manuscript. (page11, line237-240)

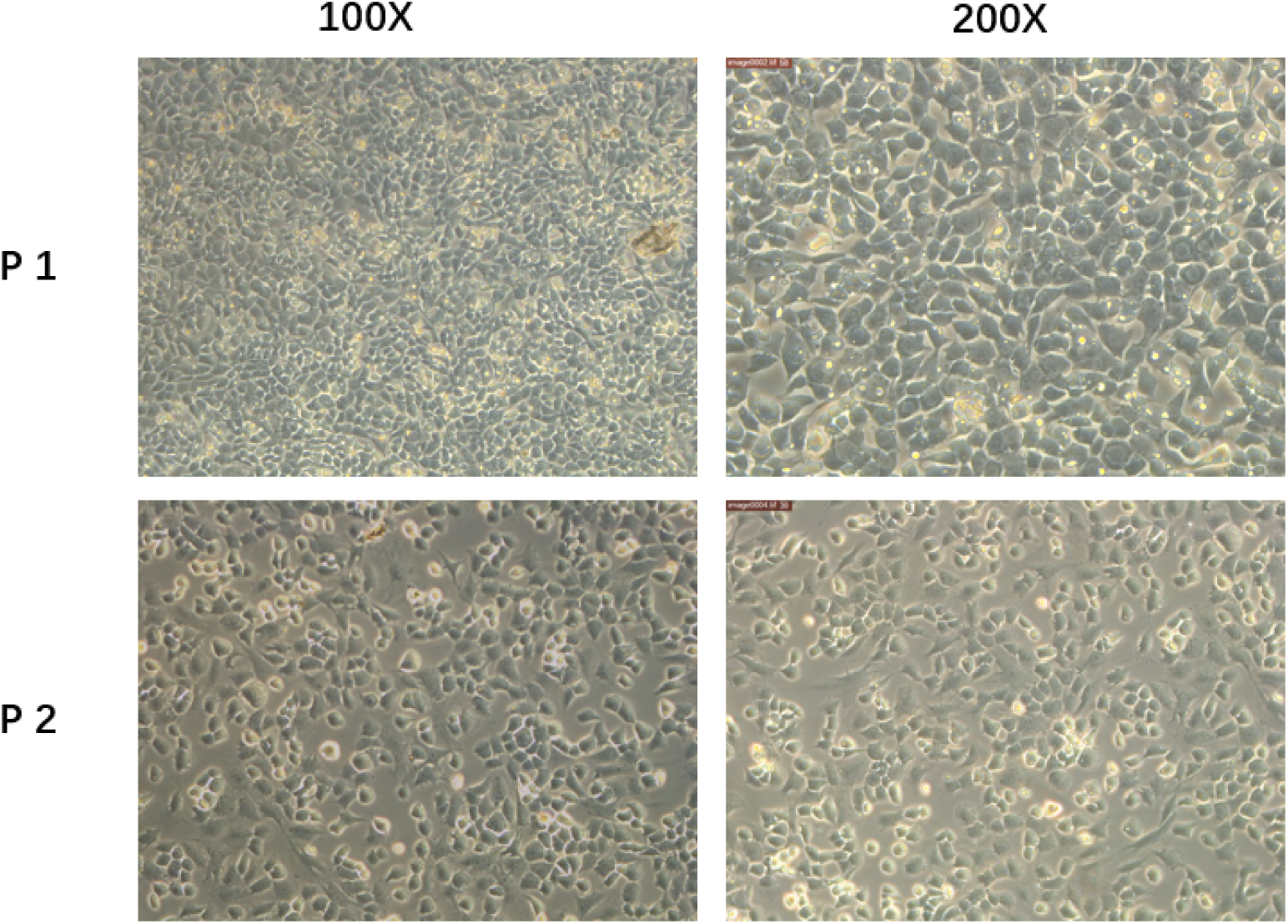
Primary mice chondrocytes we cultured (P1)and the secondary generation cells(P2) we used in the following experiment.

**References which used 2D** :

3. Liu Q, Zhai L, Han M, et al. SH2 Domain-Containing Phosphatase 2 Inhibition Attenuates Osteoarthritis by Maintaining Homeostasis of Cartilage Metabolism via the Docking Protein 1/Uridine Phosphorylase 1/Uridine Cascade. Arthritis & Rheumatology

5. Page 7, lines 148-149: “The cartilage of mice injected with the miR-199b-5p mimic was slightly degraded (p=0.02) (Fig. 4E, F)”. However, there was no significance between the groups found in Fig. 4F. Also, from the histological images of Fig. 4E, it looks like mice with inhibitor injection had more cartilage damage than miR-199b-5p mimic.

***<u>Response:</u>***We apologize for any confusion caused. Figures 4E and 4F represent the Safranin Fast Green Staining staining of the joint after the administration of miR-199b-5p inhibitor and mimic under physiological conditions. As you can see, there is minimal difference between these four images. There is no statistically significant difference. However, in Figures 5E and 5F, the MIA-induced KOA model was utilized, and noticeable differences can be observed after the administration of the inhibitor and mimic. In the revised version, we have emphasized that Figures 4E and 4F represent the results under physiological conditions, not under the MIA-induced model. (page 7, line 146-151)

6. Page 7, lines 149-150: “Additionally, the articular surface showed insect erosion (Fig. 4G).” It is also unclear how micro-CT analysis will be able to demonstrate the erosion of cartilage. Or the authors actually indicate the trochlear groove. However, this could also be observed in the control group and the results were not quantified. It is also unclear if the cross-section images of micro-CT shown here are helpful at all without any further explanation in the manuscript.

***<u>Response:</u>***Figure 4 G represents control, vehicle control, inhibitor, and mimic groups, while Figure 5 G represents model, model+vehicle control, model+inhibitor, and model+mimic groups. From Figure 4G, it can be observed that the simulator group showed the most obvious erosion appearance, while the inhibitor group did not exhibit this phenomenon^5^. From Figure 5G, it can be seen that the model group and model+mimic group exhibited the most pronounced erosion appearance, while the model+inhibitor group showed the best recovery. To highlight the pathological changes in the erosion appearance, we marked the typical locations with red arrows in the images for easy comparison and reading by the readers (Fig. 4G; Fig. 5G). We also made corresponding textual modifications in the original manuscript to address these findings (page 7, line 150-151; page 8, line 160-161). In addition, the 3D reconstruction of micro-CT is based on the synthesis of these cross-sectional images.

**References**

5. Tao Y, Wang Z, Wang L, et al. Downregulation of miR-106b attenuates inflammatory responses and joint damage in collagen-induced arthritis. Rheumatology (Oxford,

7. Page 17, line 309-310: “Before model establishment and at 3, 7, 10, 14, 21, and 28 days after model establishment.” Please re-write this as this is not clear regarding the experimental procedure.

***<u>Response:</u>***Thank you. We had to re-write the sentences as following:Baseline testing of behavioral pain thresholds was conducted prior to model establishment, followed by behavioral pain threshold testing on days 3, 7, 10, 14, 21, and 28 after model establishment. (pages15, line322-324)

8. Fig. 5A. The M + inhibitor and Model images are not at the same plane as M + mimic and M + RNAnc images.

***<u>Response:</u>***Thank you. We have modified.

9. Fig. 5B. There are two lines both with circle markers (Control and M+inhibitor). Please correct.

***<u>Response:</u>***We have corrected.

10. Fig. 5F. Missing * sign.

***<u>Response:</u>***We added *sign.

11. Please elaborate how the potential binding sites between miR-199b-5p and Gcnt2 and between miR-199b-5p and Fzd6.

***<u>Response:</u>***We apologize for any lack of clarity in the original text. In fact, we utilized targets to predict potential binding sites. Specifically, for the mouse species, we predicted that the 3’UTR of *Fzd6* binds with miR-199b-5p at positions 2483-2490, 3244-3251, 3303-3309, and 3854-3860, while the 3’UTR of Gcnt2 binds with miR-199b-5p at positions 2755-2762 and 4144-4151. In the revised version, we provide a detailed description of the methodology used for predicting these sites and offer an elaborate explanation of the results. (pages16, line352)

Additionally, to demonstrate consistency with human binding sites, we not only predicted the binding sites of human miR with these two target genes but also found a high conservation of up to 95.65% between the human and mouse sequences of miR-199b-5p. We have included this information in the supplementary materials (Fig. S6).

In Fig. 6E-F, we presented the potential binding sites between miR-199b-5p and Gcnt2, as well as between miR-199b-5p and Fzd6. In addition, we provide the predicted binding of human sequence to illustrate the binding sites. Furthermore, the predicted binding of human miR-199b-5p with *fzd6* and *gcnt2* showed a high degree of consistency. (The fluorescent labeling in the following text indicates the potential predicted binding sites.) (Supplement file 8)

hsa-miR-199b-5p MIMAT0000263

CCCAGUGUUUAGACUAUCUGUUC

NCBI Gene ID 8323 GenBank Accession NM_001164615

Gene Symbol FZD6 3’ UTR Length 1368

Gene Description frizzled class receptor 6

3’ UTR Sequence :

agaacattttctctcgttactcagaagcaaatttgtgttacactggaagtgacctatgcactgttttgtaagaatcactgttacattcttcttttgcacttaaagttgcattgcctactgttatactggaaaaaatagagttcaagaataatatgactcatttcacacaaaggttaatgacaacaatatacctgaaaacagaaatgtgcaggttaataatatttttttaatagtgtgggaggacagagttagaggaatcttccttttctatttatgaagattctactcttggtaagagtattttaagatgtactatgctattttacttttttgatataaaatcaagatatttctttgctgaagtatttaaatcttatccttgtatctttttatacatatttgaaaataagcttatatgtatttgaacttttttgaaatcctattcaagtatttttatcatgctattgtgatattttagcactttggtagcttttacactgaatttctaagaaaattgtaaaatagtcttcttttatactgtaaaaaaagatataccaaaaagtcttataataggaatttaactttaaaaacccacttattgataccttaccatctaaaatgtgtgatttttatagtctcgttttaggaatttcacagatctaaattatgtaactgaaataaggtgcttactcaaagagtgtccactattgattgtattatgctgctcactgatccttctgcatatttaaaataaaatgtcctaaagggttagtagacaaaatgttagtcttttgtatattaggccaagtgcaattgacttcccttttttaatgtttcatgaccacccattgattgtattataaccacttacagttgcttatattttttgttttaacttttgttttttaacatttagaatattacattttgtattatacagtacctttctcagacattttgtagaattcatttcggcagctcactaggattttgctgaacattaaaaagtgtgatagcgatattagtgccaatcaaatggaaaaaaggtagttttaataaacaagacacaacgtttttatacaacatactttaaaatattaaggagttttcttaattttgtttcctattaagtattattctttgggcaagattttctgatgcttttgattttctctcaatttagcatttgcttttggtttttttctctatttagcattctgttaaggcacaaaaactatgtactgtatgggaaatgttgtaaatattaccttttccacattttaaacagacaactttgaatacaaaaactttgttttgtgtgatcttttcattaataaaattatctttgtataagaaaaaaaaaaaaaa

hsa-miR-199b-5p MIMAT0000263

CCCAGUGUUUAGACUAUCUGUUC

NCBI Gene ID 2651 GenBank Accession NM_001491

Gene Symbol GCNT2 3’ UTR Length 2780

Gene Description glucosaminyl (N-acetyl) transferase 2 (I blood group) 3’ UTR Sequence:

gctattcatgagctactcatgactgaagggaaactgcagctgggaagaggagcctgtttttgtgagagacttttgccttcgtaatgttaaccgtttcaggaccacgtttatagcttcaggacctggctacgtaattatacttaaaatatccactggacactgtgaaatacactaacaggatggctgggtagagcaatctgggcactttggccaattttagtcttgctgtttcttgatgctcacctctatattagtttattgttaggatcaatgataaatttaaatgacctcagatctttgcaccagatactcatcatatacaaatgttttagtaaaaaagagaattgtagataatactgtctaggaaaataagaattaggtttctttgaagaaggaatcttttataacaccttaacagtcaccactgtgctcaaccagacagatagtgaaacagctttctgggtaattcaccaatttcctttaaaacataagctacctgaatggagaatacatcttgtttctgagtttcaacactagcatttttggcttactcatggacaaagttctgtatatagtataaagtcattaacaagaaaca ggatatgctttaagacagaattcactgtctgttgcttcagtaaaaggacctcggggaataaaacatttctctcttatatgccagaatgtaggctggtccctatgtcatgtcttccattaagaacactaaaaagtccttgcaagaatggagatatgcattcaagagaggtgctatcacatagatctagtctgaagtctggaacactttcctcttctatgacccctctctccccagtattatcttacttgcaaaatggagaccaaattctatcctgtgaggcttttaattgcaccatagtatgctctgagtagctttacactgcctggtactgatagtagtggctcgatttttaagagccttcaattgtagatgaacatctctgttatttatccctcattcatccatccgttcattcattcagccttcaatcaacatctcttgagtgtctattatgtacaggacatgtactgagacaaaaaggaaacataagagctttttcactctaaaaatcttggcaataatgtcaacaccagaaagcctcctctggagaatcttacagagtgattgtagtttaatacaggaacacacagggctgtgtagcatgataccaggcccaggagatcagtaattacaaattaagggttaaatcagagattattcaacagagagggagaaaggaggagacagagggaggacctgttgtgttccagccattctggtattcctttatgtatctaatttcattcaaacctcacaacagtcttgtgaggcccttatataattactcccattttgcagatgaagtaactgaggcttagaaaggttaatagcaccggggaacaatttctctgggtgagaattgggactctgttgctggtcttctcagttcatttcctgaggtggatttactgagagaaggtgaaataaagccatatttagtataccagagaaggtagattttaagaatggtctcagtgttaatactgagaaaaagtcctgtcagttcagaaaaaatgtgaagtctactttagtattcctgtaatactaaaccgttgagtttctaaatatttatttattctaacaaaaagcaattactacaaatggatgacacatttaatgaacacaattttattttttttctgtaactgtgcttgttgaatgtcaatcatatttaaagggaatgactttgaagtaaaaccttttttcttgctactgaaaaaaatggagttgttttgggtggtaaagtgttaaggaatagggacagctggtcacacaaggaactcttgaaggccacatgtgaaaacctgtcacttgcacagaggccagtcccactaaggtgaccagagtgggctccaagcacaaactgccattggctatagatgggactgtgtccccccaaaattcatgtgttggagccttaaccctcaatgtgatggtatttgagatggggcctttggtaagggaagtttagatgaggtcacgagggtaggaccctcatgatgggatgagtccccttacaagacctctggcttgggccgggcgtggtggctcacacctgtaatcccaacactttgggaggccaaggcaggtagatcacttgatgccaggagttccagaccaggctggccgacatggtgaaaccccatctctactaaaaaatataaaaattagccgggctttgtggcatgtgcctgtaatcccagctatttggcaggctgaggcatgagaatcgcttgaacccaggaggtggaggttacagtgagctgagagtgccccactgcactccagcctgggtgacagagcgagactttgtcccaaaacaaaataggtgaggggatagcgaatgcactcagggtcagcagtggagtttaaaaattgtctcttttcaacttatttaaatgacagcacctgagaagaggaaccgttttacactggatgtttctcatgtagaacaagaaatctttctggaattgatgtttacatgtctgttgttggtcatctctcctgtgtcttaaatactttaatgttggaagagcatagtgtttgggctagtgggtttctgacagcccatgggaatgccctgaaactactgtatctgatgtttgttttcgatgaggttccatgttttgttttcttgggaataaattaatatattgttttccaaaaaaaaaaaaaaaaaaaa

12. Page 10-11, Line 222-223: “Our findings indicate that miR-199b-5p plays a crucial role in KOA by targeting Fzd6 and Gcnt2”. This is an overstatement. The current work shows the possible bindings of miR-199b-5p and Fzd6 as well as bindings of miR-199b-5p and Gcnnt2. Whether miR-199b-5p truly functions through Fzd6 and/or Gcnt2 requires genetic knockdown of Fzd6 and Gcnt2 in the presence of miR-199b-5p. Thus, please tune down this statement and the title of the manuscript.

***<u>Response:</u>***We agree your opinion of our conclusion. Therefore, we delete the overstatement sentences and tune down the conclusion of the manuscript. (the title; page 8,179; page11, line227-228)

13. The Schematic figure (the last figure). Please remove osteophyte as this was not quantified in the study.

***<u>Response:</u>***We modified the schematic figure accordingly.

Minor concerns:

1. Most figures were distorted.

***<u>Response:</u>***We provide a new version of the figure to avoid distortions.

2. Providing GO term numbers in Fig. 1C is not very helpful. Maybe show the GO term and corresponding numbers in the manuscript (Page 4, lines 79 - 82).

***<u>Response:</u>***Thank you for your advice. We added the corresponding notes of the GO term numbers in the manuscript to explain each biological concept of it. (Page 4, line 77-89; Page 22, line 515-532)

3. What were M-0.5 and M-1 in Fig. 2D? Different MIA concentrations?

***<u>Response:</u>***Yes, these are different MIA concentrations, which we illustrate in the legend. (Page 23, line 535-536)

4. Please follow the nomenclature of the gene symbol. For example, Fig. 3E-P should be mouse genes (?).

***<u>Response:</u>***We modified the relevant gene symbol.

5. Page 3, line 59. Not all chondrocytes are pathogenic cells in OA.

***<u>Response:</u>***we are sorry for the mistake, now it has been modified. (Page 3, line 59)

6. Typo. Page 3, line 55.

***<u>Response:</u>***We changed the Typo.

7. Page 4, line 78. These are differentially expressed miRNAs, not genes.

***<u>Response:</u>***We have revised the unsuitable expression. (Page4, line75-76)

I wish the authors all the best with their continued work in this area.

***<u>Response:</u>***Thank you for your wishes.

## Notes

### Competing Interest Statement

The authors have declared no competing interest.

