## Supplementary figures and images for "Inhibition of miR-199b-5p reduces pathological alterations in Osteoarthritis by potentially targeting *Fzd6* and *Gcnt2*"

### supplement fig1

supplement fig 1

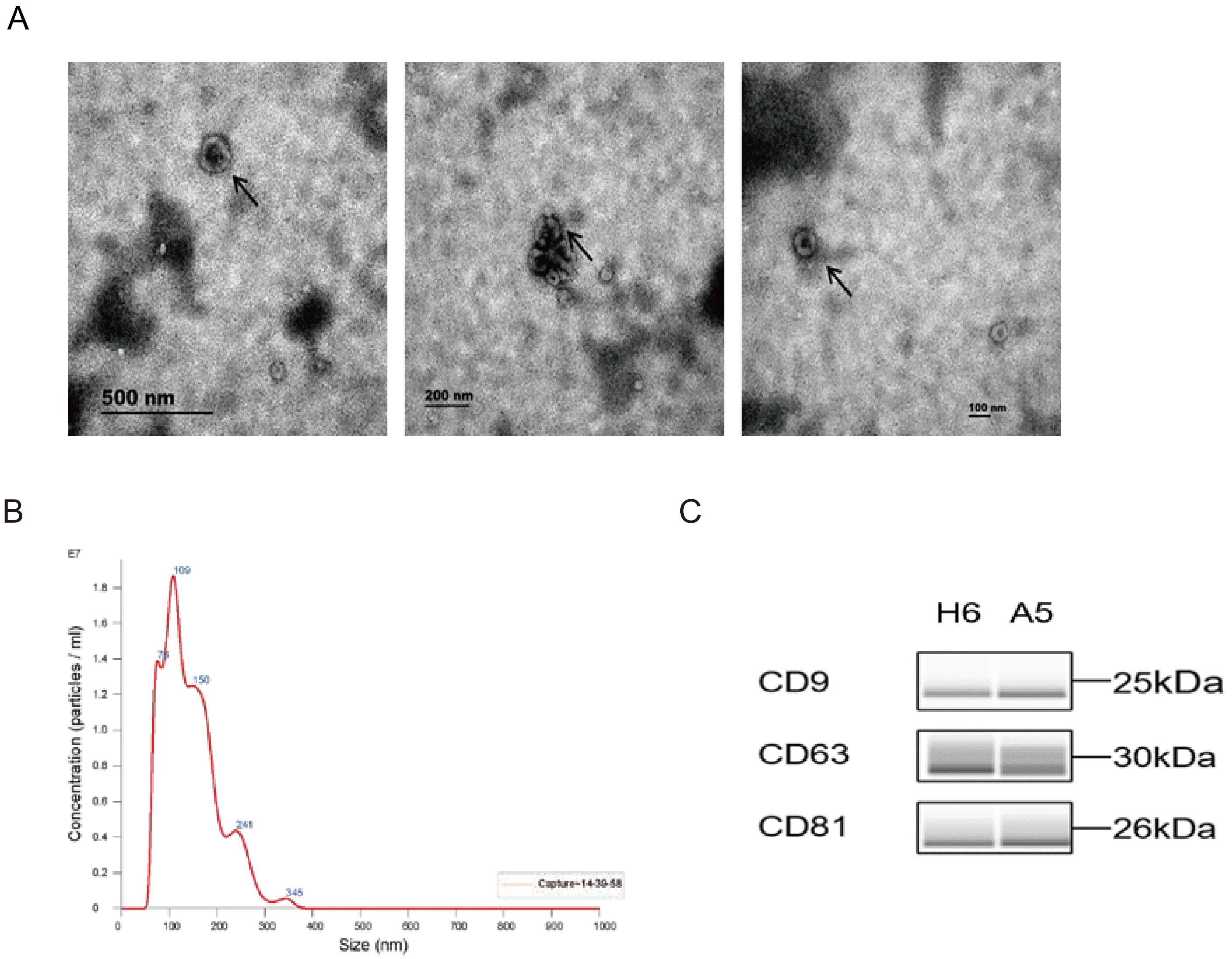

### supplement fig2

supplement fig2

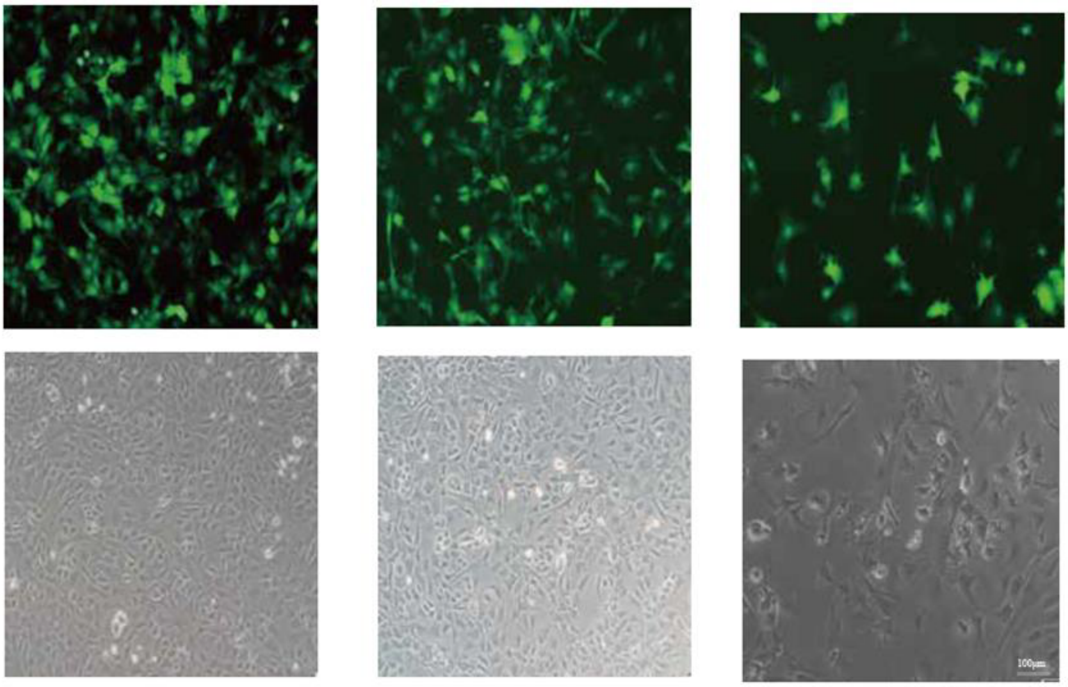


RNAnc inhibitor mimic

### supplement fig3

supplement fig 3

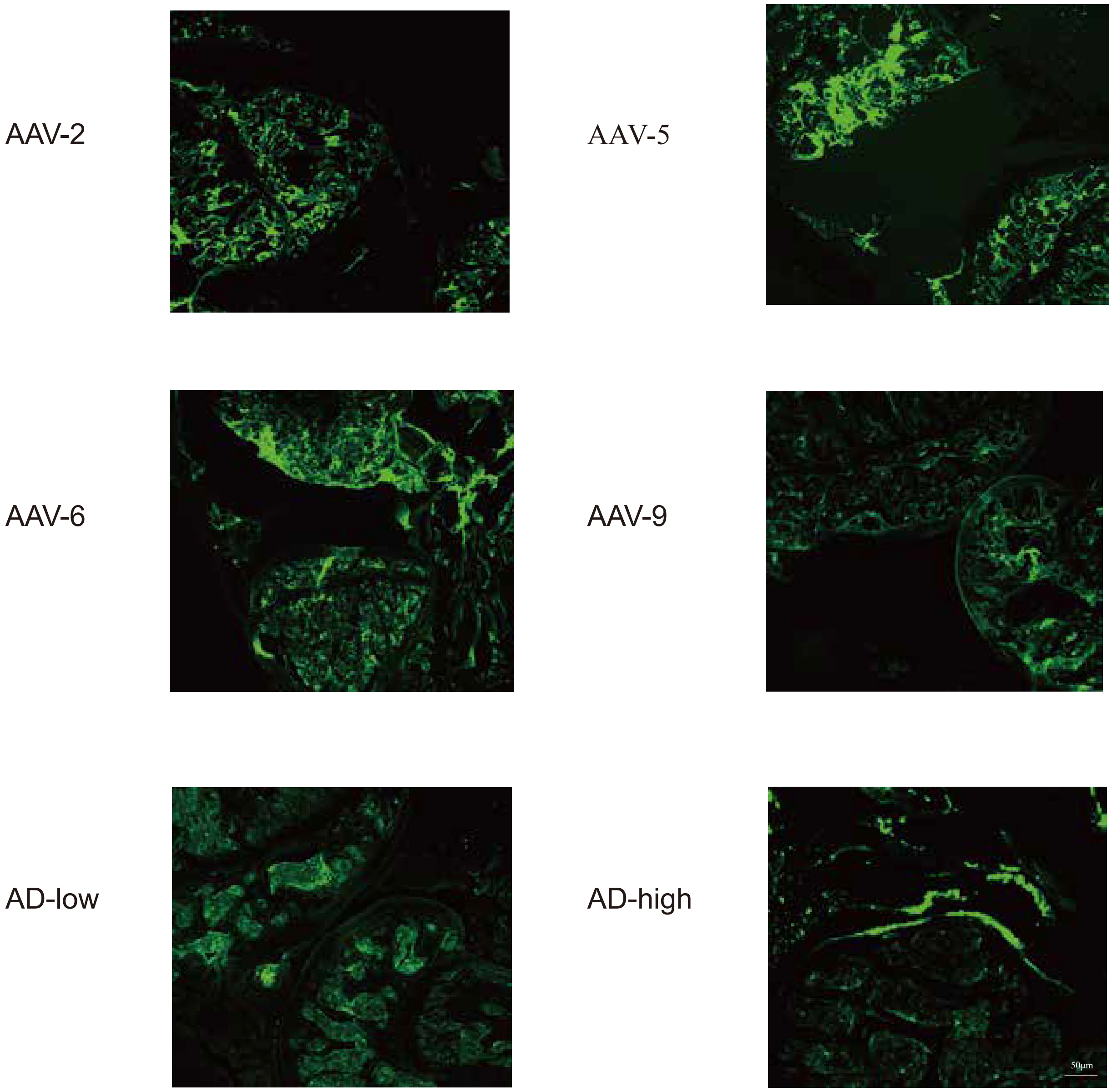

### supplement fig4

supplement fig 4

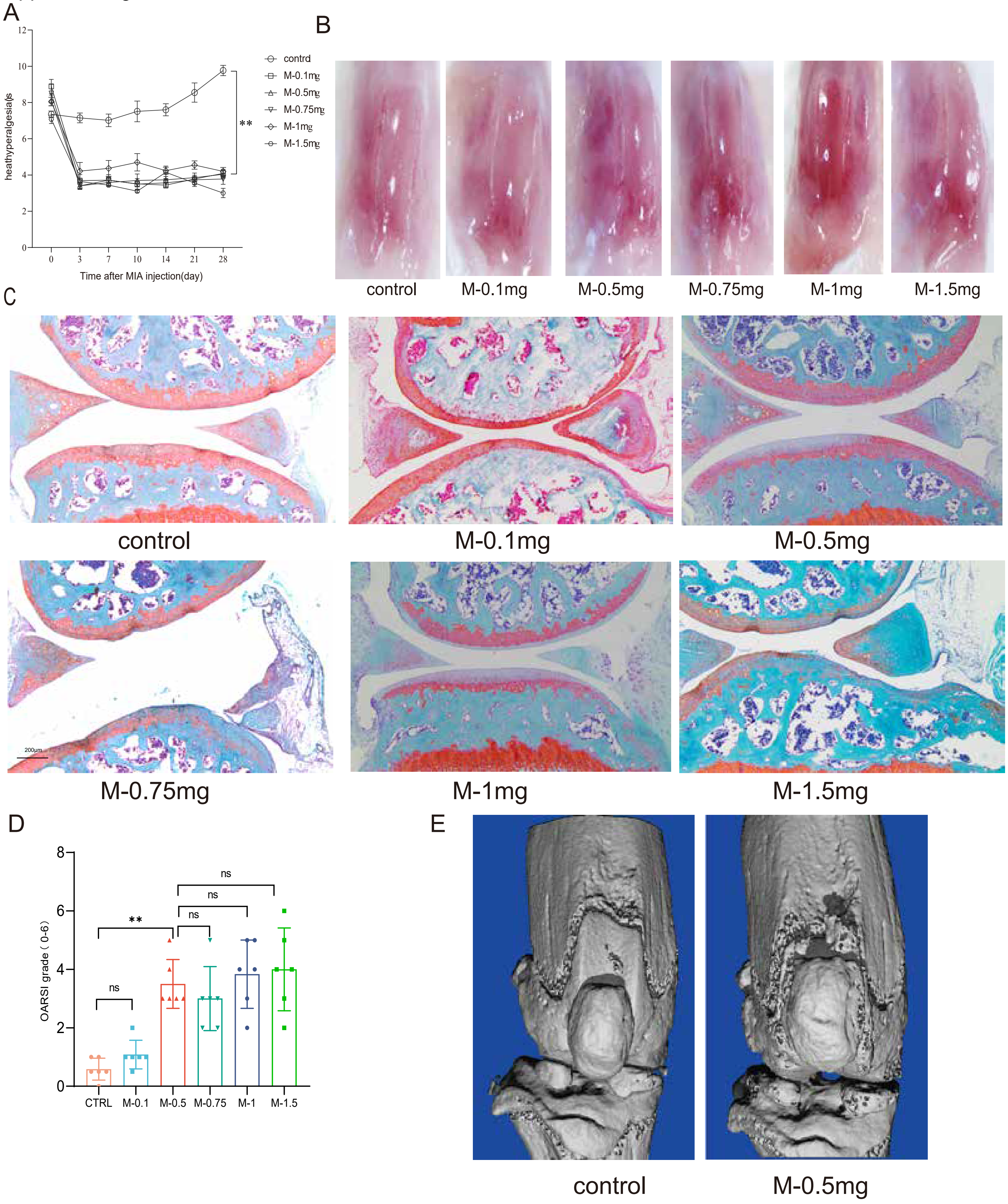

### supplement fig5

supplement fig 5

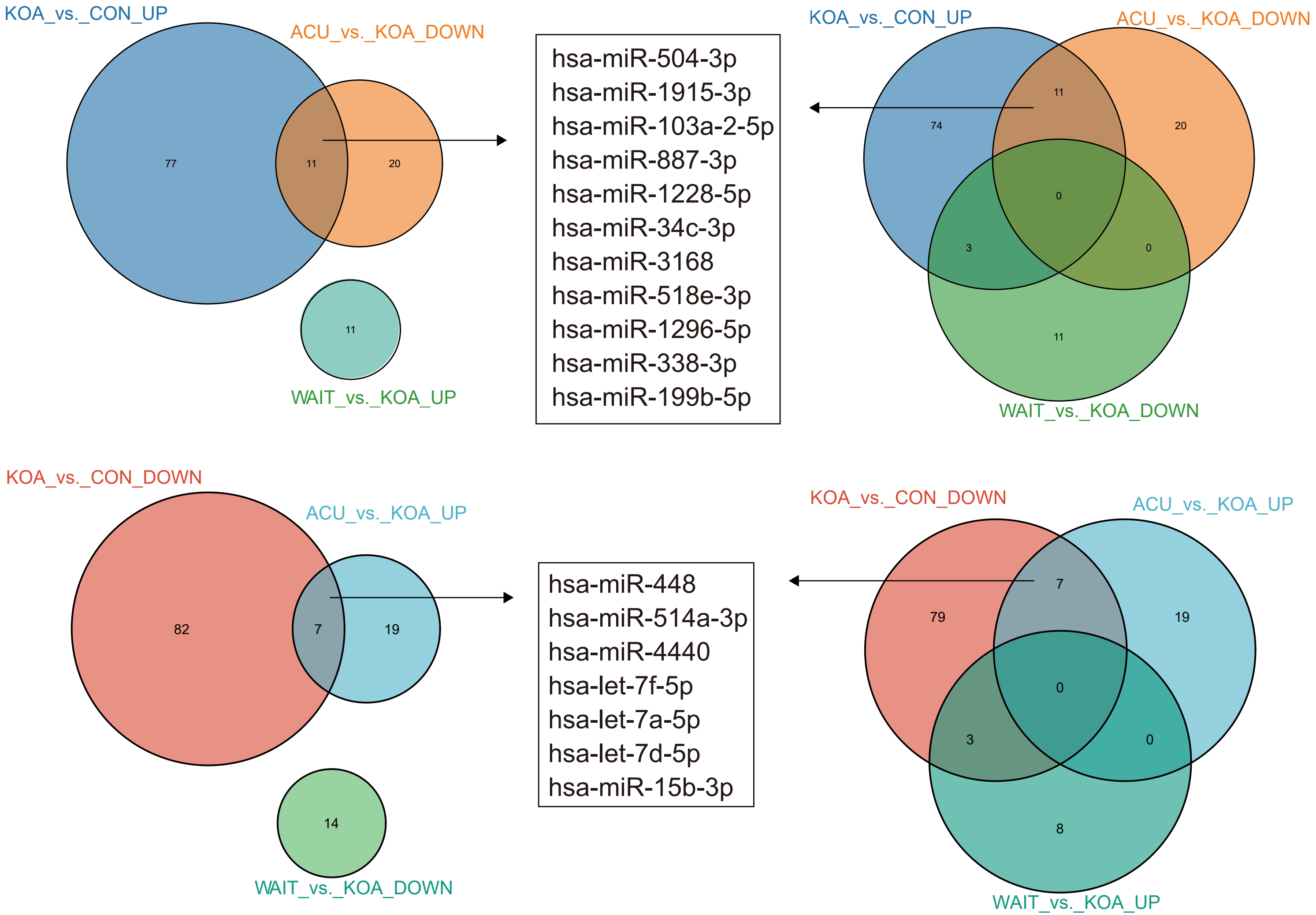

### supplement fig7

supplement fig 7

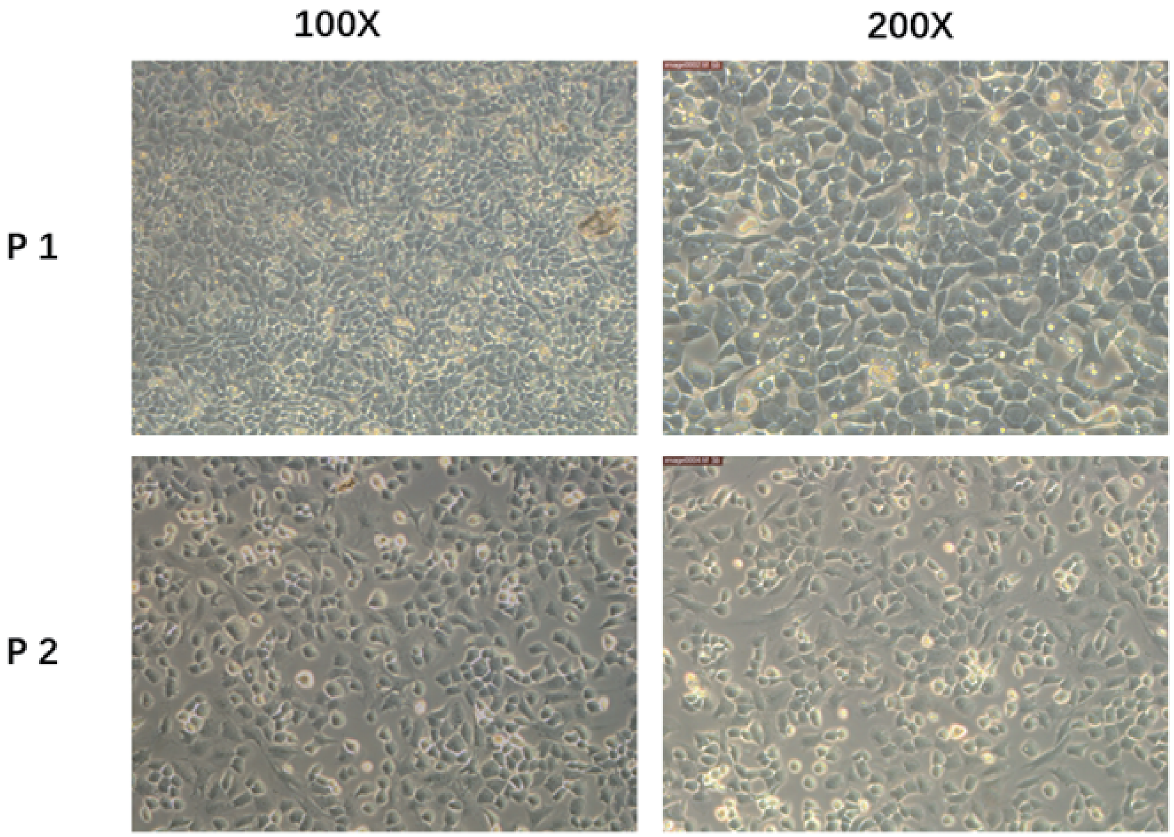
