## supplement fig6 for "Inhibition of miR-199b-5p reduces pathological alterations in Osteoarthritis by potentially targeting *Fzd6* and *Gcnt2*"

**supplement fig 6**

A

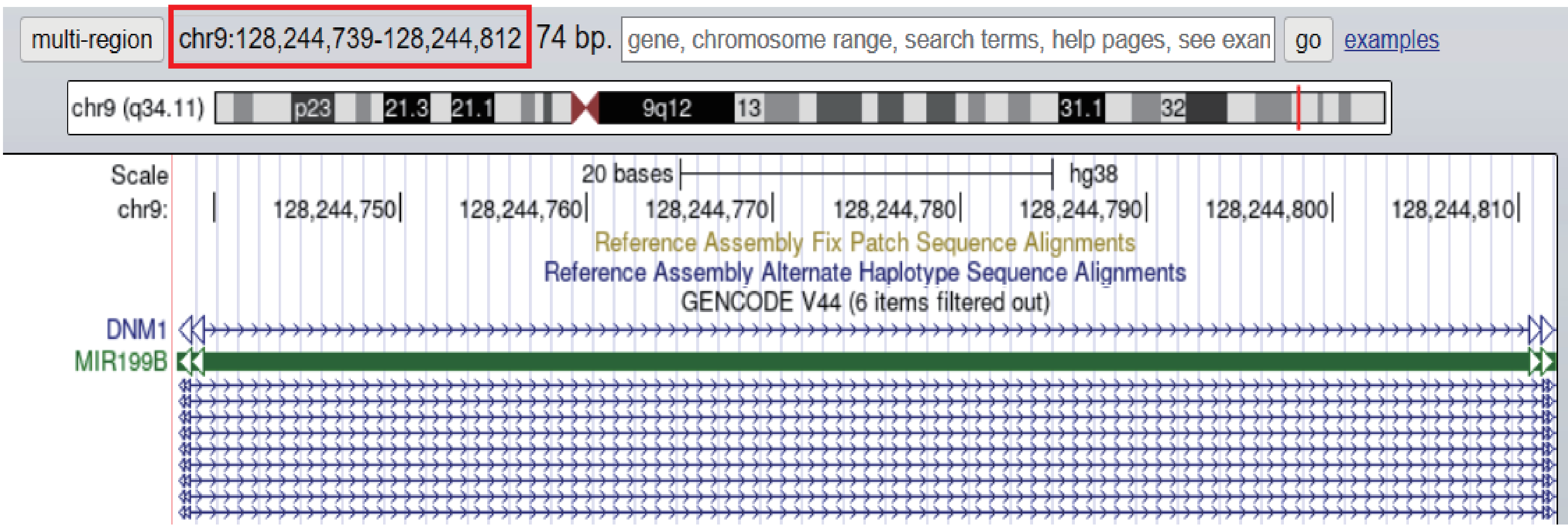


B

| | |
|---|---|
| pre–miRNA | hsa–mir–199b miRBase |
| Genomic Coordinates | chr9: 128244721 – 128244830 |
| Mature miRNA | hsa–miR–199b–5p |
| Sequence | 26\| CCCAGUGUUUAGACUAUCUGUUC \|48 |

C

>mmu–miR–199b–5p MIMAT0000672
CCCAGUGUUUAGACUACCUGUUC

hsa-miR-199b-5p MIMAT0000263
CCCAGUGUUUAGACUAUCUGUUC

D

**Sequences producing significant alignments** Download

Select columns

Show 100

select all 1 sequences selected Graphics MSA Viewer

| | Description | Scientific Name | Max Score | Total Score | Query Cover | E value | Per. Ident | Acc. Len | Accession |
|---|---|---|---|---|---|---|---|---|---|
| ☑ | hsa-miR-199b-5p | | 38.2 | 38.2 | 100% | 1e-09 | 95.65% | 23 | Query_111461 |
