## supplement file8 for "Inhibition of miR-199b-5p reduces pathological alterations in Osteoarthritis by potentially targeting *Fzd6* and *Gcnt2*"

>mmu-miR-199b-5p MIMAT0000672

CCCAGUGUUUAGACUACCUGUUC

UGUGACC ACACTGG

UGUGAC	ACACTG

GUGACC	CACTGG

Mus musculus frizzled class receptor 6 (Fzd6), transcript variant 1, mRNA

NCBI Reference Sequence: NM_008056.3

AGAAAACTGTCTCGTTCCCCCAGAAGCACATGTATGTTACACTGGAGATGACCAACTGATTTGTCTTATAAAGGCCACTGTTGAGCTGGGAGAGTAGCCCAGTGGTACAGCGCCCACCTGGAATACTTGAGGACCTGGGGTTGTCTCCCAGCACTGCAAAAGGAAAATTCACTGTTACAGTCTTCCTTGCACTTAAACCAGCTTTGTCTATTGTTTTTTTGGTTTGGCTTTTATTTTTGTTGCTGTTATTTTTGTTGTTGTTTGTTTGTTTTTTTGTTTGTTTGTTTGAGACAGGGTTTCTTTGCTAGCCCTGACTGTCCTGAAACTCCCTCTGTAGACCAGGCTGGCCTCAAACTTACAGAGATCTGCCTGCCTCAGCCTCCCGAGTGCTGGGAATAATGGTGTGGTCACCACTGCCCAGCCTTTTGTCTGTTTTTAAACTTGAAAGAAACAACAGCCCAGATTTCAAAAATAATATAATGCATTTATACCTAAAAAACCAACCAGGAGTGCCCAGTTAATAACACTTTTTAAATGTGGGGATGGGAAGGGCATTAGAGGAGTCTTCCTTCTATTGAAGATTCATTAAAGTATTTTAAGATATGCTCTTTCACTCTTTATATAAATCCAAGATTTTTCTTTGCTGAAGTATTTAAAACTTTTGTACCTTTATATGTAGATATGAATTTGAAAATATGCTTATGTGTATTTGAACTTTTGAAAATCCTAGAGAATTGAATCAAATATTTTTATGATGTTTTTCTACTATTTTAGCTACTTTGCGACTGTGATAGCTGTTACACTGGATTTTTAAAAAACTTGTACAGCAGCCTCTTTACAGTAAAAAGAGTGGGTGTCACACTGAAAGGTCTGTAAGAAGTGGTCACAGCCACCCCTACCTTCCCCAAAAGGAGGAACTTGGTGGCAGGTCCCTCCCTGATTGGACTGTCCCTTTCTTTCTGCATGTTATAAATCAGCAGGTAAGATGGTAGGTTTTTACAAGTTAGGCCGAGCTGTCGATTCCCCTTTTAAGTGTTGAATTAGGATTGAATTATGGCCATTTGTAGTTGCTCGTGCCTGTCTTTATTTTAGTATTTTATTTCCCGAGACAGGAACTCACTGTGTGGTGCTCCTTGGCTGTCTGGTGTTCAGTCTGTCCCAGGCAGGTCACAGAGATCTCCCCCTCTGCAGCCCACTCATCTCTCCCAAGCCACCACACTCAGCTTTTATCTGTTTTAAAAATTTAAACTTAAAAAAATGTTTTTGGAATAGTACAAACACATTGTGTTGTAAATTTCTTTGATGCTATGCAAAATTCCTATCTGCATCTAAGCCTGCAAAAGAAAATGTGCGAAGGGCAGAGTCAGAGTTGGGCAGGAAGAGTGTAGTGCAGCAGATGCAGCGTGAAGACACTGAAGGTGCTAAGACAGCGTCTCAGTGCTGGTCCTCCTTAAGGATTATCTCGCCAGCGAGGTTTTCTTAGATACTTTGATCCCATTGGAGCTCTGTTAAAGTTTAAAATGAAAATTATCATGTACTGTATGGGAAATGTAAATACTAACTTTTCCACATATGTAAACTTCAGACACAAATTTTTTTGTGTGTTCTTTTCATCAATAAAATTTTCTTTGTATAAAAAAA

拼接序列：

AGAAAACTGTCTCGTTCCCCCAGAAGCACATGTATGTTACACTGGAGATGACCAACTGATTTGTCTTATAAAGGCCACTGTTGAGCTGGGAGAGTAGCCCAGTGGTACAGCGCCCACCTGGAATACTTGAGGACCTGGGGTTGAATTGAATCAAATATTTTTATGATGTTTTTCTACTATTTTAGCTACTTTGCGACTGTGATAGCTGTTACACTGGATTTTTAAAAAACTTGTACAGCAGCCTCTTTACAGTAAAAAGAGTGGGTGTCACACTGAAAGGTCTGTAAGAAGTGGTCACAGCCACCCCTACCTTCCCCAAAAGGAGGAACTTGGTGGCAGGTCCCTCCCTGATTGGACTGTCCTATCTGCATCTAAGCCTGCAAAAGAAAATGTGCGAAGGGCAGAGTCAGAGTTGGGCAGGAAGAGTGTAGTGCAGCAGATGCAGCGTGAAGACACTGAAGGTGCTAAGACAGCGTCTCAGTGCTGGTCCTCCTTAAGGA

>hsa-miR-199b-5p	MIMAT0000263

CCCAGUGUUUAGACUAUCUGUUC

NCBI Gene ID	8323	GenBank Accession	NM_001164615

Gene Symbol	FZD6	3' UTR Length	1368

Gene Description	frizzled class receptor 6

3' UTR Sequence ：agaacattttctctcgttactcagaagcaaatttgtgttacactggaagtgacctatgcactgttttgtaagaatcactgttacattcttcttttgcacttaaagttgcattgcctactgttatactggaaaaaatagagttcaagaataatatgactcatttcacacaaaggttaatgacaacaatatacctgaaaacagaaatgtgcaggttaataatatttttttaatagtgtgggaggacagagttagaggaatcttccttttctatttatgaagattctactcttggtaagagtattttaagatgtactatgctattttacttttttgatataaaatcaagatatttctttgctgaagtatttaaatcttatccttgtatctttttatacatatttgaaaataagcttatatgtatttgaacttttttgaaatcctattcaagtatttttatcatgctattgtgatattttagcactttggtagcttttacactgaatttctaagaaaattgtaaaatagtcttcttttatactgtaaaaaaagatataccaaaaagtcttataataggaatttaactttaaaaacccacttattgataccttaccatctaaaatgtgtgatttttatagtctcgttttaggaatttcacagatctaaattatgtaactgaaataaggtgcttactcaaagagtgtccactattgattgtattatgctgctcactgatccttctgcatatttaaaataaaatgtcctaaagggttagtagacaaaatgttagtcttttgtatattaggccaagtgcaattgacttcccttttttaatgtttcatgaccacccattgattgtattataaccacttacagttgcttatattttttgttttaacttttgttttttaacatttagaatattacattttgtattatacagtacctttctcagacattttgtagaattcatttcggcagctcactaggattttgctgaacattaaaaagtgtgatagcgatattagtgccaatcaaatggaaaaaaggtagttttaataaacaagacacaacgtttttatacaacatactttaaaatattaaggagttttcttaattttgtttcctattaagtattattctttgggcaagattttctgatgcttttgattttctctcaatttagcatttgcttttggtttttttctctatttagcattctgttaaggcacaaaaactatgtactgtatgggaaatgttgtaaatattaccttttccacattttaaacagacaactttgaatacaaaaactttgttttgtgtgatcttttcattaataaaattatctttgtataagaaaaaaaaaaaaaa

>mmu-miR-199b-5p MIMAT0000672

CCCAGUGUUUAGACUACCUGUUC

UGUGACC ACACTGG

UGUGAC	ACACTG

GUGACC	CACTGG

Mus musculus glucosaminyl (N-acetyl) transferase 2, I-branching enzyme (Gcnt2), transcript variant 1, mRNA

NCBI Reference Sequence: NM_008105.3

CCCGCAGCAGCTCGGGCCTAAATGGAAATTGAAGACGTAAAGAAGAGCTTTCTTTTCCAAGAGACTCTGGTCTTGGCTATGCTGAAGACTTTTTTAAAAAATGGTTTTCAGGGAACCGTGAGGATCTGGCAACATGGCTCTGCTTGCAATATCCACTGAGCACTGTAATACATTTGACAGGATGGCTGCATGAAACTGTCCAGCATTTTTCTGTCGTTTCCCCCCCACCCCAAATCCTTTTTTTCTCTTCTTTTTTAAAAGATAGGTAGGGTCTTGCTAGATATCTCTAGCAAGCCTGTTTCTTGTCAGGTAGCCCAGGATGGCCTTGAACTCTTAATCCTCCGCCTTGGCCCTCTAAGGGCTGCTAGTCCGAGTGTGGACAACCACACCTGGCTCTGGCGTCTGGCGGTCACTTGACGCTCATCTGCGTCTCCGTCTGCTAGGTGGTCGGTGATAAATTCAAATGTATCAGATCTCTGTGCCAGATACTCATATTAGTATACCCTGTTCAAAGGAGAATGGTGGATAATACTGTCTGAGGAAATCAGAACTTATTTGAGAAGGAAGCTCTCGTTCCACATTCCTAGCAACTATATTCTCAGGCACACATAGGCGCACAGAGACAGCTTCCTAGTAACTCACATCAATTTCGTTTAATACAAACTGATCCTGAAGAGCTGAGGAACTATGTAAGTTCAGTGTCCAAGCATTTGACTGACATTTGTGCAGAGTTAGGTTCCATCTCCAGCATCACGCACGGGTCTTGTGAACACCGACATCTTCAGTTTACTTGTGTCTGCAGTTCTGTGGACAGAACAAAGCCATTCACAAGTGACCGACAGCACGTGCCGTGAGACGGGAGGTGGCTGCAGGAAGAGGATGCCTCAGGGATGCCGGACATTTCCGTGTTCTATTCCAGAGTGTTCTCTTCTCCCAGAGAACACTGGAACGTCCTCACAGGGATGAAGGAATTCTTTCTTTCCCACAGGAAAGGTGCTACTGTGTGGGTCTGGAACCTGTGAATGAATCACTCCTGGGGTTTCGTTTCTATGTGAGAGACGGGGACCCGGTTCTGTTGTGTGCAGCTTTAGCGGCATCAGGACAGCTTCCAGCTTGCAATGATTGCTGCTGACAGGAACACCTGGATCTCAAGAGCTTTTTATGACAGGTGGACATCTGTGTCATTTTTCCTTGTTCCCTCATCCATCCATCCATCCATCCATCCATCCATCCATCCATCCATCCATTCGTCAGCCTACATTCAACAAACGCCTCTTGAATGTCTATGATTCCCCCAGACACATGCTGACAGACACGAGGGAAGCAAGACAGATTCTTTCTCCACTGTTAAACTCTGGGGAGTGGTATCAGCTTTTGAAACCCTGCTCTAGACTCTTACAAAATGGCTACACTTAGCACATACTCTGCCCTGGCTATATATTATATTGGGCCATAGAGATCAGCAATGTATCAATTAATAGATTAATCCAGAGACAGGAAGGAGGAAACGAAAGCTAGAGCCTCTTTGTGACTCCACCAACTCCTTCTTGTCTCCTTTTATTGAAGAGTCCGCACACATACTCACCCCTCTGGTGGGCCACATGAGGACCTGCATCTTGCAGATGATGTAGCAAGCCTTAGGATGGTTCATGATCAGTAGGAAGCAAATTCTAAGACCTCAGGGTGTCTTTCTGAGTCTCTCATTTCTTTAGGGATGTGGAGAGAAGATGAAGTATAGCCGTGTGTAGTGTCGCCAGAGAAGGTGGAGTTTAAGAATGGTCTCAGTGTTATTAATGAGAAAAAGTTCTGTCATTCAGGAAAAAAAAGCAAAAAACGTGAGGTTGGGACTCAGTAGGTGAGGGTGCTTGCAGCCAAACCTGATGACCTGCATTCAATTCCTGGGTCCCACGGGACAGAGGGAGAGAGTCCACTACTGAAAGGAACGCTCAAACCTCCTCACACAGGTCCTTGTGCCTGGGCACTCCCCCGCCCCATATGCATACACACATACATGCTTTTGTGATTTTTTTTATTAGTGCTTTTACATTTAGAAATATTTATTTATTCTAGCAGGGAAAAAACCCCACATGCTTTAAATGATGTCACTTTCCACAACTTCATTCTTGTGCTGTCATTGGACTGAATGTTATGCATCTTTAAAGGGAATCACTTCGTGTAAAACTCTCTTGTCTCTTCTGCTGAAAAGTAAAATGGGTTGTGTCGGGTAATAAAGGGTCGAGGGGTAGGAAGCACTTGGTCAAGGCGAAGTCTAAAACCGGTCTCTTTTTTATTTGTTACTTGGATGAGAGCACCTGAAGACTAATGTTTTACACTGGATTGAGATCTCATGGAAAATGAAACATTCTTTTTCTGGAGAAGTTTACATGCCCATTGTTGGTCACCTCTTTACTGTAAATACTATTGAAAAAAGGAGCGTAGCATTTGCATAGCAGGGATTGTGCCTGTATCTACTGTGTCCATTTTGTTTTTTCGGATGTTTTCTGGTTTTGTTTCCTTGGAAATAAATTAATATATTGTTTTCCACCTTCAAAAAAAAAAAAAAAAAAA

拼接序列：

AGTTCTGTGGACAGAACAAAGCCATTCACAAGTGACCGACAGCACGTGCCGTGAGACGGGAGGTGGCTGCAGGAAGAGGATGCCTCAGGGATGCCGGACATTTCCGTGTTCTATTCCAGAGTGTTCTCTTCTCCCAGAGAACACTGGAACGTCCTCACAGGGATGAAGGAATTCTTTCTTTCCCACAGGAAAGGTGCTACTGTGTGGGTCTGGAAAACTCTCTTGTCTCTTCTGCTGAAAAGTAAAATGGGTTGTGTCGGGTAATAAAGGGTCGAGGGGTAGGAAGCACTTGGTCAAGGCGAAGTCTAAAACCGGTCTCTTTTTTATTTGTTACTTGGATGAGAGCACCTGAAGACTAATGTTTTACACTGGATTGAGATCTCATGGAAAATGAAACATTCTTTTTCTGGAGAAGTTTACATGCCCATTGTTGGTCACCTCTTTACTGTAAATACTATTGAAAAAAGGAGCGTAGCATTTGCATAGCAGGGATTGTGCCT
